# miRNA-encoded peptide, miPEP858, regulates plant growth and development in Arabidopsis

**DOI:** 10.1101/642561

**Authors:** Ashish Sharma, Poorwa Kamal Badola, Chitra Bhatia, Deepika Sharma, Prabodh Kumar Trivedi

## Abstract

MicroRNAs (miRNAs), small non-coding endogenous RNAs, are processed product of primary miRNAs (pri-miRNAs) and regulate target gene expression. pri-miRNAs have also been reported to encode small peptides, miRNA-Encoded Peptides (miPEPs). Though regulatory role of miPEPs has been speculated, no detailed study has been carried out to elucidate their function through developing knock-out mutants. Here, we report that pri-miR858a of *Arabidopsis thaliana* encodes a small peptide (miPEP858a) which regulates the expression of pri-miR858a leading to modulation in the expression of target genes involved in the plant growth and development as well as phenylpropanoid pathway. CRISPR-based miPEP858a-edited plants developed phenotypes similar to that of mature miR858-edited plants suggesting crucial role of miPEP858a in mediating miR585a function. miPEP858a-edited and miPEP858a overexpressing lines altered plant development and accumulated modulated levels of flavonoids due to changes in expression of associated genes. Exogenous treatment of synthetic-miPEP858a to the miPEP858a-edited plants complemented phenotypes and the gene function suggesting a significant role of miPEP858a in controlling the miR858 function and plant development.

**One sentence summary:** Small peptide, miPEP858a, encoded by primary miRNA for miR858a regulates plant growth, development and flavonoid biosynthesis

The authors responsible for distribution of materials integral to the findings presented in this article in accordance with the policy described in the Instructions for Authors

## Introduction

MicroRNAs (miRNAs) are small non-coding, single-stranded endogenous RNAs which regulate gene expression mainly through cleavage and/or translational repression of target mRNA in a sequence-specific manner (Catalanotto et al., 2016; Vasudevan, 2012; Ambros, 2001; Pillai, 2005; Chen, 2009; Bartel and Chen, 2004; Song et al.,2011; Nithin et al., 2015). These miRNAs act as the key regulators of a variety of plant and animal developmental processes (Liu et al., 2018; Yu et al., 2017; Reinhart et al., 2002; Jones-Rhoades et al., 2006). Biogenesis of miRNA requires well-coordinated multiple crucial steps (Bartel, 2004; Kurihara and Watanabe, 2004). miRNA gene is first transcribed by RNA Pol II to form a primary miRNA (pri-miRNA), which is usually several hundred nucleotides long (Bartel, 2004; Lee et al., 2004). The pri-miRNA is further processed to generate a stem-loop like intermediate called precursor miRNA or pre-miRNA, by the Drosha RNase III endonuclease in animals (Bartel, 2004; Lee et al., 2002) or by Dicer-like 1 enzyme (DCL1) and HYPONASTIC LEAVES1(HYL1) and SERRATE (SE) proteins in plants (Kurihara and Watanabe, 2004; Tang et al., 2003). Subsequently, these pre-miRNAs are processed leading to mature miRNAs which play important role in the regulation of gene expression. Over the past few years, a number of studies have demonstrated the role of miRNAs as vital regulators of plant development processes, nutrient uptake as well as stress responses through targeting mostly transcription factors (TFs) (Kumar et al., 2017; Tang and Chu, 2017).

Recently, it has been demonstrated that the plant pri-miRNA contains short Open Reading Frames (ORFs) which encode regulatory peptides also known as miRNA-Encoded Peptides (miPEPs) (Lauressergues et al., 2015).These peptides were shown to enhance the mature miRNA levels by enhancing the transcription of their associated pri-miRNA (Couzigou et al., 2015; Couzigou et al., 2016). These miPEPs show specificity towards associated miRNAs suggesting their wide application in the regulation in gene expression and improvement of the desired agronomically important traits (Couzigou et al., 2017). Though there are only a few reports on involvement of miPEPs in the gene regulation, these provide a significant glimpse into the importance of these small peptides.

Flavonoids synthesized by the phenylpropanoid pathway, are secondary metabolites produced in plants and are reported to have role in pigmentation, as UV-protectants, signaling molecule and many other biological functions (Winkel-Shirley, 2001;Buer and Muday, 2004;Santelia et al., 2008; Quattrocchio et al., 2006; Stracke et al., 2007; Misra et al., 2010; Seleem et al., 2017). Recently, we demonstrated that the miR858 plays a crucial role in the development of Arabidopsis and regulation of the phenylpropanoid pathway.miR858 has been reported to down-regulate the expression of various MYB transcription factors post-transcriptionally, which includes MYB11, MYB12 and MYB111 (Sharma et al., 2016; Wang et al., 2016). MYB11, MYB12, and MYB111 have been demonstrated to regulate the biosynthesis of flavonols by targeting genes that encode early pathway enzymes including Chalcone synthase (CHS), Chalcone isomerase (CHI), flavonoid 3 hydroxylase (F3H), and flavonol synthase (FLS) (Pandey et al., 2014;Mehrtenset al., 2005; Stracke et al., 2010).

Here, we show that pri-miR858a of *Arabidopsis thaliana* encodes a small peptide (miPEP858a) which regulates the expression of miR858a at the transcriptional level and thereby affects the expression of target genes involved in the phenylpropanoid pathway as well as plant growth and development. Exogenous application of synthetic miPEP858a resulted in enhanced miR858a expression along with substantial down-regulation of associated target genes like flavonol-specific *MYBs* with marked changes in plant growth and development. To provide a better insight into the miPEP858a function, for the first time, we edited the coding region of miPEP858a as well as miR858 family members using the CRISPR/Cas9 system. miPEP858a-edited plants showed phenotypes similar to that of mature miR858-edited plants. These phenotypes include reduced plant growth and delayed flowering along with the enhanced accumulation of flavonoids, anthocyanin and reduction in the level of lignin accumulation.Constitutive expression of miPEP858a showed modulated expression of miR858a and target genes as well as contrasting phenotype as compared to miPEP858a-edited plants. Moreover, to strengthen miPEP function, we used synthetic miPEP858a for the complementation of CRISPR-edited miPEP858a and GUS reporter studies. Our results demonstrated that the exogenous treatment of synthetic miPEP to these plants suggest a significantly important role of miPEP858a in controlling and regulating the miRNA function.

## Result

### Effect of miPEP858a on phenotype in Arabidopsis seedlings

As a first step, to identify miPEP encoded by pri-miR858a, we screened for the putative open reading frames (ORFs) in 1000 bp region upstream from pre-miR858a in *Arabidopsis thaliana* (Col-0) (Supplemental Figure 1). The analysis led to the identification of 3 putative ORFs of 156 bp, 135 bp and 39 bp respectively. To study whether these ORFs are active *in planta,* constructs were developed for the in-fusion expression of β-glucoronidase (GUS) reporter gene with translation initiation codons (ATG^1^, ATG^2^ and ATG^3^ respectively) of three identified ORFs along with upstream promoter region. Transient expression of these constructs in *Nicotiana benthamiana* leaves and histochemical GUS staining showed that leaves infiltrated with construct carrying ATG^1^ of the ORF of 135 bp (ATG^1^) was active *in planta* conditions (Figure 1A). These results suggest existence of 135 bp long ORF encoding small peptide (miPEP858a) of 44 amino acid residues upstream of pre-miR858a region (Figure 1B). To study whether this peptide is translated in plants, a construct was prepared in which the complete ORF^1^ along with upstream promoter region was fused to GUS and transiently expressed in *Nicotiana benthamiana* leaves. This result demonstrated that ORF^1^ is translated *in planta* and might therefore regulate miR858a activity (Figure 1A).

**Figure 1.**
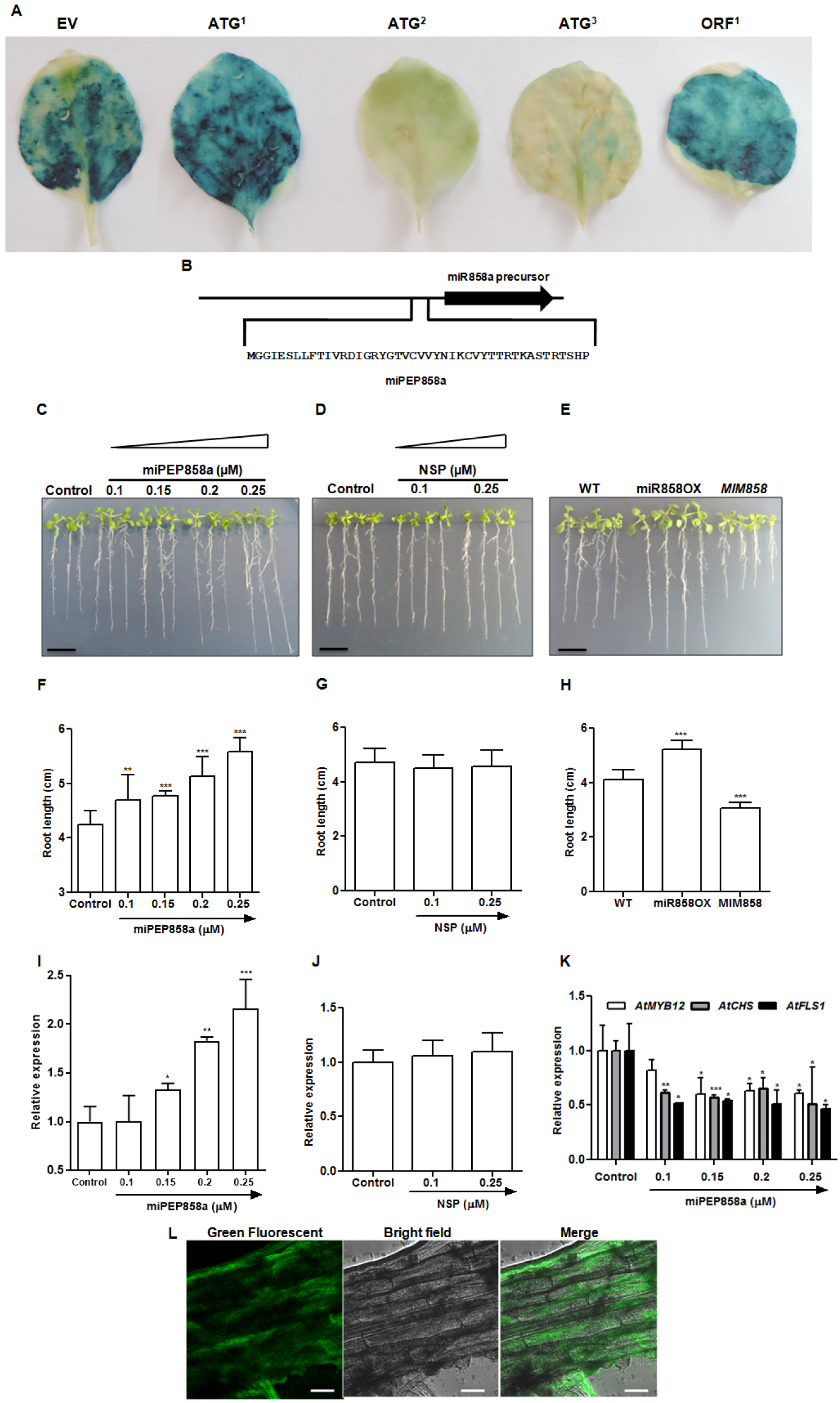
Effect of miPEP858a on Arabidopsis seedlings. (**A)** GUS assay in *Nicotiana benthamiana* leave with EV, Pro858:ATG^1^:GUS, Pro858:ATG^2^ :GUS, Pro858:ATG^3^ :GUS and Pro858:ORF^1^:GUS. (**B)** miPEP858a sequence (44 amino acid residues) encoded by pri-miR858a. (**C)** Representative photographs of the phenotype of 10-day old WT (Col-0) seedlings grown on ½-strength Murashige and Skoog (MS) medium supplemented with water (control) or with different concentrations of miPEP858a. (Scale bars, 1 cm). (**D)** Phenotype of 10-day old WT seedlings grown on ½ MS supplemented with water (control) or two different concentrations of non-specific peptide (NSP) (scale bars, 1 cm). (**E)** Phenotype of WT, miR858OX and MIM858 transgenic lines grown on ½ MS medium for 10 days (scale bars, 1 cm). (**F)** The root length of 10-day old WT supplemented with water (control) or with different concentrations of miPEP858a. **(G)** The root length of 10-day old WT supplemented with water (control) or two different concentrations of non-specific peptide (NSP). (**H)** Root length of 10-day old WT, miR858OX and MIM858 transgenic lines. (**I, J)** Quantification of pre-miR858a in WT seedlings treated with water (control), different concentrations of miPEP858a and non-specific peptide (NSP). (**K)** Quantification of miR858a target genes in WT seedlings treated with water (control), different concentrations of miPEP858a peptides. Data are plotted as means ±SD. Error bars represent standard deviation. Asterisks indicate a significant difference between the treatment and the control according to two-tailed Student’s t-test using GraphPad Prism 5.01 software (n=30 independent seedlings, * P < 0.1; ** P < 0.01; *** P < 0.001). (**L)** Green fluorescence, Bright-field (DIC) and merged microscopic images of Arabidopsis root cells in the presence with 50μM 5′FAM-miPEP858a after incubation for 12 h. Scale bar = 50 µm

As miPEPs modulate accumulation of mature miRNA by enhancing transcription of its corresponding pri-miRNA (Lauressergues et al., 2015), we hypothesized that the application of miPEP858a might affect miR858a expression and associated phenotypes. To test whether miPEP858a is functional, WT seeds were grown on the plant growth media supplemented with synthetic peptide miPEP58a at various concentrations (0.1 to 0.5 μM). The root length of seedlings grown on media supplemented with exogenous miPEP858a showed a concentration-dependent increase as compared to growth in control media (Figure 1C and F). This increase in root length is comparable to the phenotype observed in seedlings over-expressing miR858a (Figure 1E and H). Based on preliminary dose-dependent experiments, 0.25 μM concentration was selected as optimal for the regulation of miR858a.

In our earlier study, miR858a was shown to target R2R3 family of MYB transcription factors including flavonol-specific activators MYB11, MYB111 and MYB12 (Sharma et al., 2016). These MYBs are known to positively regulate flavonol biosynthesis genes such as *CHS, CHI, F3H* and *FLS1* (Stracke et al., 2008). We performed qRT-PCR analysis to investigate the role of miPEP858a on the expression of pri-miR858a and target genes (MYB transcription factors and genes associated with flavonoid biosynthesis) (Sharma et al., 2016; Rosany et al., 2018; Piya et al., 2017). Results suggested enhanced transcript levels of miR858a and substantial decrease in transcript levels of target and associated genes, in agreement with a positive role of the miPEP858a in regulating miRNA858a expression (Figure 1I and K).

In previous studies, it was shown that miPEPs are specific for their associated miRNA. Therefore, specificity of miPEP858a was studied by using a NSP (non-specific peptide of 42 amino acid residues) at two different concentrations (0.1 μM and 0.25 μM). Seedlings grown on media containing NSP showed no change in the root growth and expression of pri-miR858a as compared to WT plants (Figure1D, G and J). Furthermore, in order to check the specificity of miPEP858a towards miR858a, expression of pre-miR858b and several other miRNAs family members was analysed in miPEP858a treated seedlings was performed. This analysis revealed no significant changes in the expression of these miRNAs in response to miPEP858a supplemental Figure 2). Together, these results suggest that pri-miRNA858a encodes a small peptide (miPEP858a) which has a potential role in the regulation of miR858 expression. To analyse whether miPEP858a is absorbed by the root, fluorescent labeled miPEP858a (5-FAM-miPEP858a) was used. Results suggest that incubation of seedlings with 5-FAM-miPEP858a, leads to green fluorescence inside root cells of Arabidopsis and the uptake of this peptide by the roots (Figure 1L)

**Figure 2.**
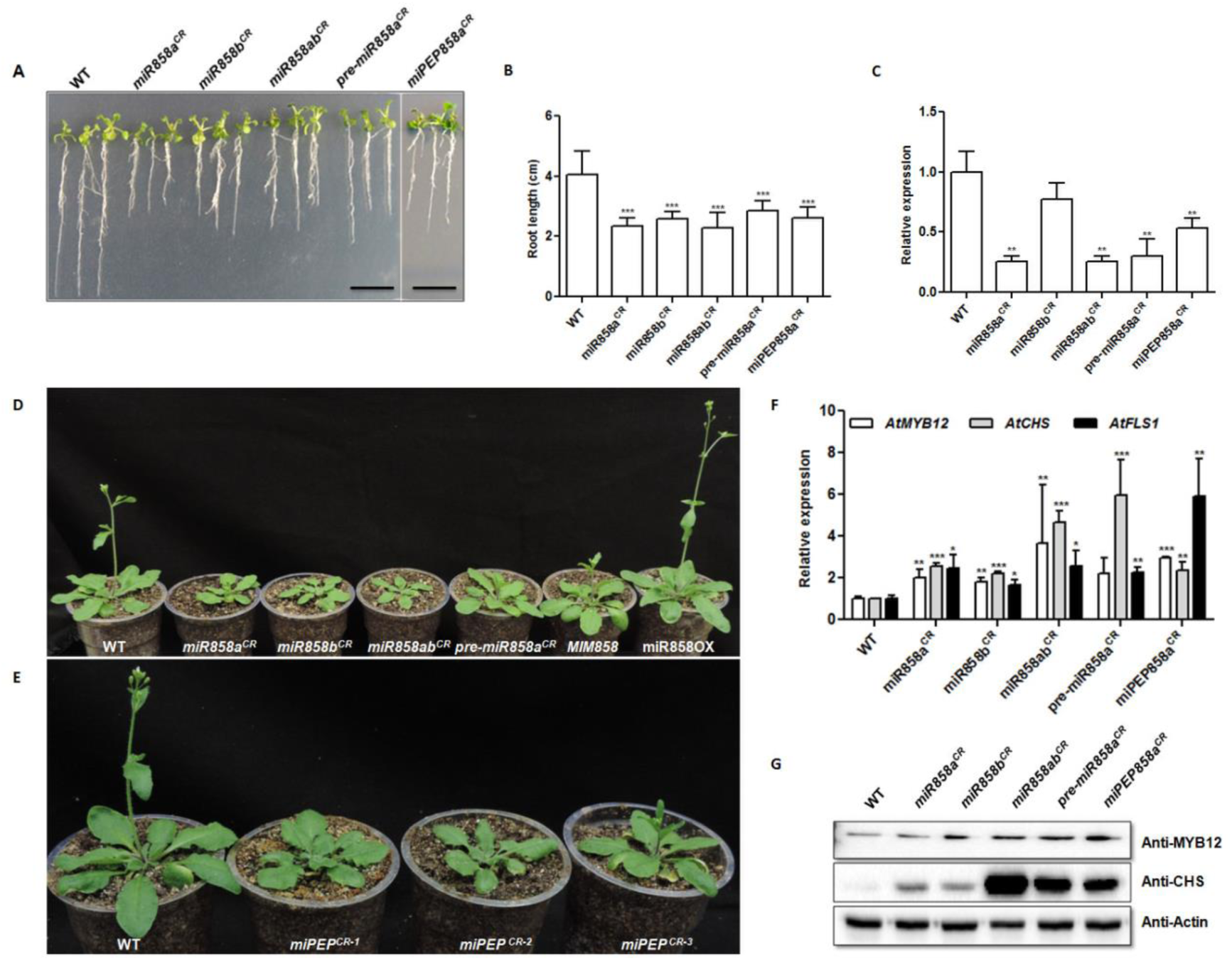
CRISPR/Cas9 derived knockout mutants show altered phenotype and gene expression. (**A)** Representative photographs of the phenotype of 10-day old WT, CRISPR/Cas9 edited *miR858a*^*CR*^, *miR858b*^*CR*^, *miR858ab*^*CR*^, *pre-miR858a*^*CR*^, *miPEP858a*^*CR*^ seedlings grown on ½ strength MS medium (scale bars, 1 cm). (**B)** The root length of 10-day old WT, *miR858a*^*CR*^, *miR858b*^*CR*^, *miR858ab*^*CR*^, *pre-miR858a*^*CR*^, *miPEP858a*^*CR*^ seedlings grown on ½ MS medium (n=30 independent seedlings). (**C)** Quantification of pre-miR858a in 30-day old rosette of WT, *miR858a*^*CR*^, *miR858b*^*CR*^, *miR858ab*^*CR*^, *pre-miR858a*^*CR*^, *miPEP858a*^*CR*^. **(D, E)** Representative photographs of the phenotype of 30-day old WT, *miR858*^*CR*^, *pre-miR858a*^*CR*^, miR858OX and MIM858 plants and *miPEP858a*^*CR*^ plant. (**F)** Quantification of miR858a target genes in 30-day old rosette of WT, *miR858a*^*CR*^, *miR858b*^*CR*^, *miR858b*^*CR*^, *miR858ab*^*CR*^, *pre-miR858a*^*CR*^ and *miPEP858a*^*CR*^ plants. (**G)** Western blot analysis of MYB12 and CHS protein in 30-day old rosette of WT, *miR858a*^*CR*^, *miR858b*^*CR*^, *miR858ab*^*CR*^, *pre-miR858a*^*CR*^, *miPEP858a*^*CR*^. Actin was used as loading control. Data are plotted as means ±SD. Error bars represent standard deviation. Asterisks indicate a significant difference between the WT and the edited plants according to two-tailed Student’s t-test using Graphpad Prism 5.01 software (n=5 independent plants, * P < 0.1; ** P < 0.01; *** P < 0.001).

### CRISPR/Cas9 derived knockout mutants show altered phenotype and gene expression

To further investigate the functions of miPEP858a, we developed miPEP858a mutant plants using CRISPR/Cas9 approach. To establish CRISPR/Cas9, we first edited *AtPDS3* (AT4G14210), a significant carotenoid pathway gene that upon mutation showed photo-bleaching (albino phenotype) (Supplemental Figure 3A) which is also reported by others groups (Li et al., 2013). For mutation in miPEP858a, we designed a gRNA which can target coding region (Supplemental Figure 4). To study if miPEP^*CR*^ plants have similar effects as miR858, knock-out miR858 mutant plants were developed. As miR858 family consists of two members, miR858a and miR858b, to develop mutation in both members a single gRNA was designed. This gRNA can target both miR858a and miR858b as both members are similar in sequences except for an additional 1bp present at the 5′ end of miR858a (Supplemental Figure 4). In addition, gRNA was designed to mutate pre-miRNA region (*pre-miR858a*^*CR*^) to study whether mutations in regions other than mature miRNAs produces similar result. For the selection of miPEP858a and miR858 edited plants, screening was performed to identify mutations in the miPEP858a coding region. Deletion and insertions were observed in the coding region that caused nonsense mutations. pre-miR858a, mature miRNA sequences of miR858a (*miR858a*^*CR*^) and miR858b (*miR858b*^*CR*^) as well as those carrying mutations in both miR858a and miR858b (*miR858a/b*^*CR*^). Homozygous and Cas9-free edited plants of T4 generation were used for further study (Supplemental Figure 5, 6 and 7).

**Figure 3.**
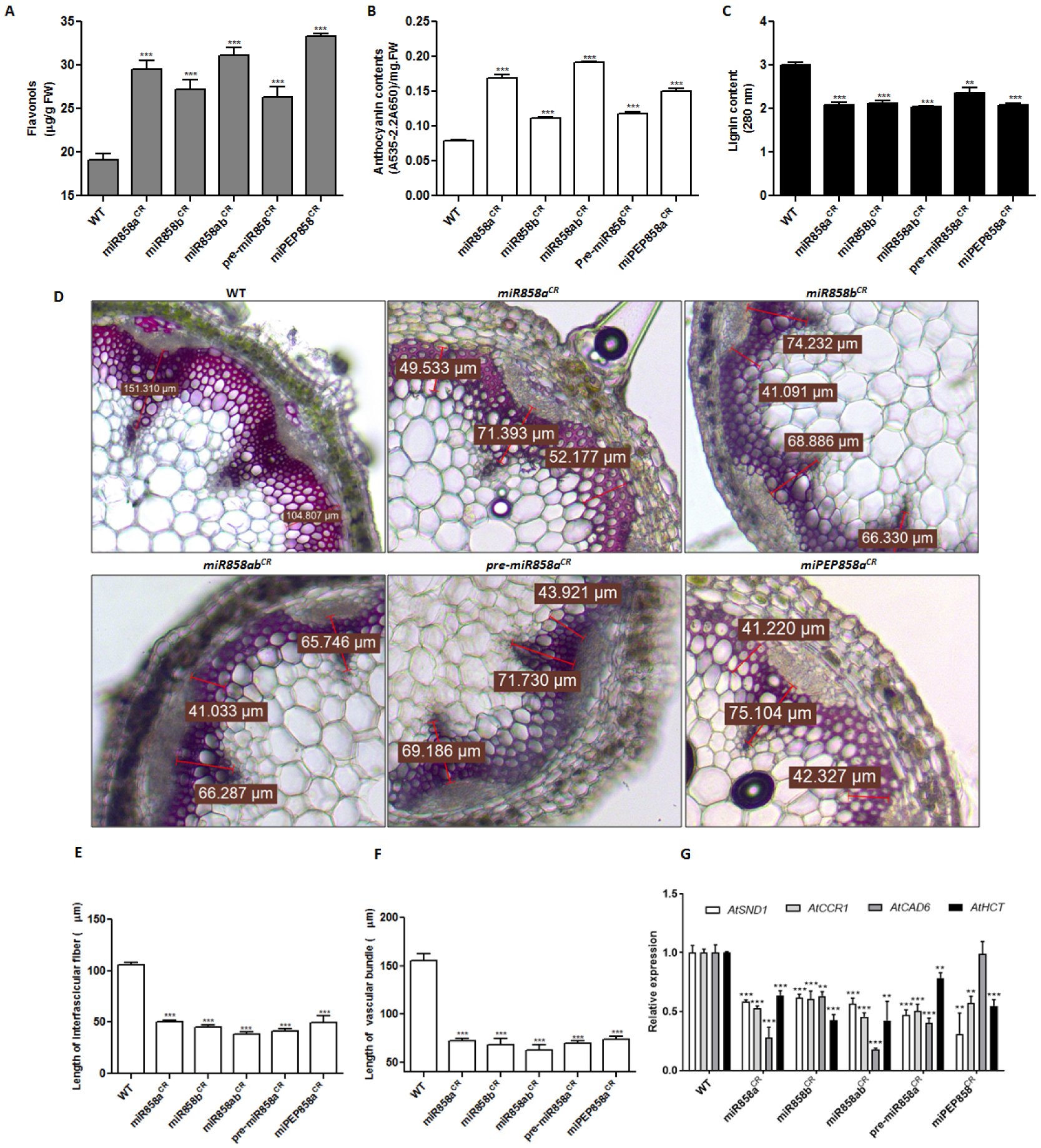
Editing of miR858 leads to differential accumulation of metabolite levels. **(A-C)** Quantification of total flavonols, anthocyanin and lignin content in 35-day old rosette and stem of WT, *miR858a*^*CR*^, *miR858b*^*CR*^, *miR858ab*^*CR*^, *pre-miR858a*^*CR*^, *miPEP858a*^*CR*^ respectively. Data are plotted as means ±SD.. Error bars represent standard deviation. (**D)** Transverse section of stem of 35-day old WT *miR858a*^*CR*^, *miR858b*^*CR*^, *miR858ab*^*CR*^, *pre-miR858a*^*CR*^ and *miPEP858a*^*CR*^ were stained with phluoroglucinol to show change in lignin content. (**E)** Bar graph showing the length of interfascicular fibers (µm) of WT, *miR858a*^*CR*^, *miR858b*^*CR*^, *miR858ab*^*CR*^, *pre-miR858a*^*CR*^ and *miPEP858a*^*CR*^ plants. (**F)** Bar graph showing the length of vascular bundles (µm) of WT, *miR858a*^*CR*^, *miR858b*^*CR*^, *miR858ab*^*CR*^, *pre-miR858a*^*CR*^ and *miPEP858a*^*CR*^ plants. **(G)** Expression analysis of lignin biosynthesis genes in WT, *miR858a*^*CR*^, *miR858b*^*CR*^, *miR858ab*^*CR*^, *pre-miR858a*^*CR*^ and *miPEP858a*^*CR*^ plants. Data are plotted as means ±SD in the figures. Error bars represent standard deviation. Asterisks indicate a significant difference between the WT and the edited plants according to two-tailed Student’s t-test using GraphpadPrism 5.01 software (n=5, independent seedlings, * P < 0.1; ** P < 0.01; *** P < 0.001).

**Figure 4.**
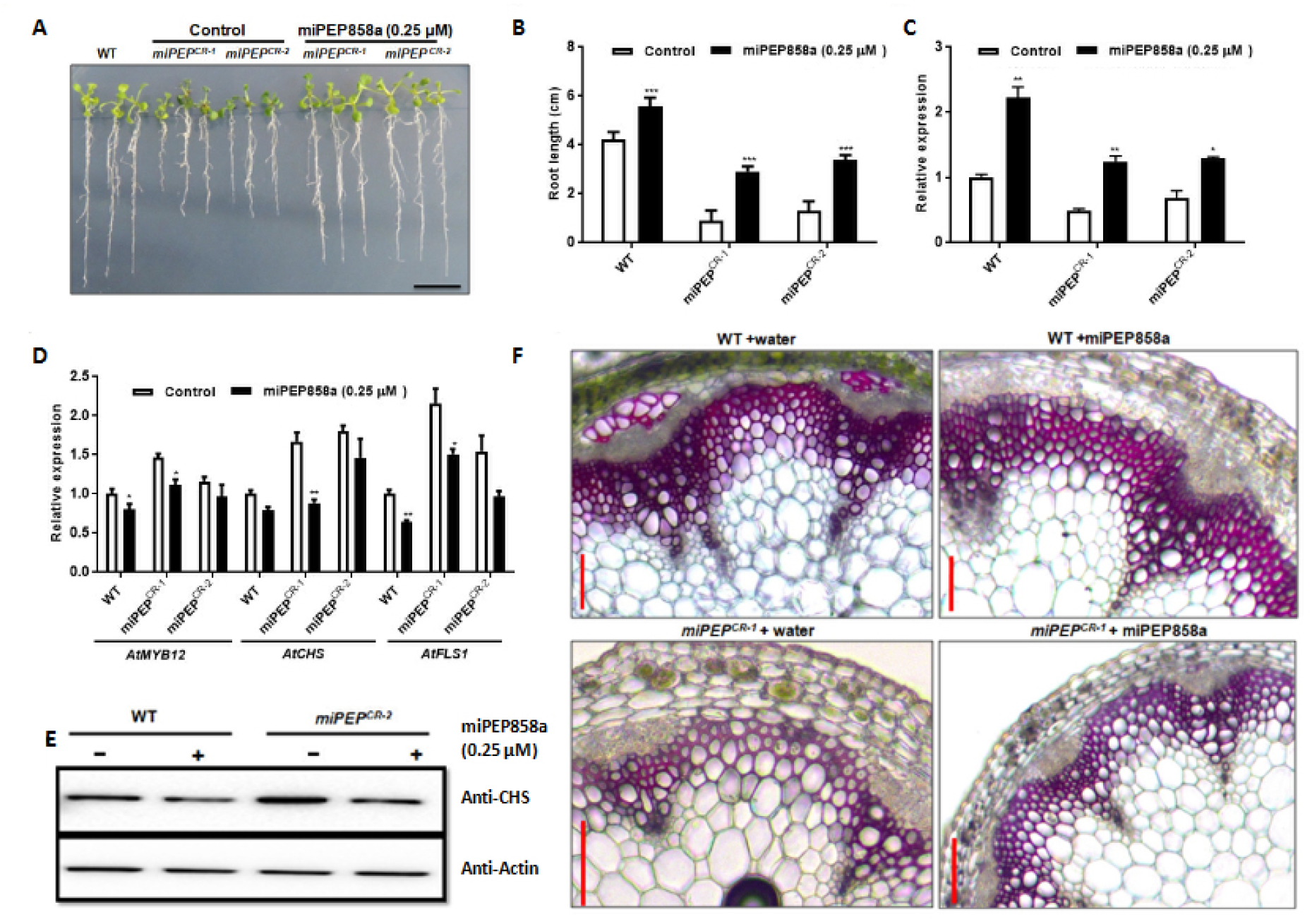
Complementation with miPEP858a synthetic peptide leads to restoration of function of CRISPR/Cas9 edited *miPEP*^*CR*^ lines. **(A)** Representative photographs of the phenotype of 10-day old WT and CRISPR/Cas9 edited *miPEP*^*CR*^ lines grown on ½ strength Murashige and Skoog (MS) medium supplemented with water (control) and 0.25 µM miPEP858a (scale bars, 1 cm). **(B)** The root length of 10-day old WT and CRISPR/Cas9 edited *miPEP*^*CR*^ lines grown on ½ strength Murashige and Skoog (MS) medium supplemented with water (control) and 0.25 µM miPEP858a (scale bars, 1 cm). (n=30 seedlings) (**C)** Quantification of pre-mir858a in 5-day old WT and CRISPR/cas9 edited *miPEP*^*CR*^ lines grown on ½ strength Murashige and Skoog (MS) medium supplemented with water (control) and 0.25 µM miPEP858a. **(D)** Quantification of miR858a target genes *AtMYB12, AtCHS* and *AtFLS1* in 5-day old WT and CRISPR/Cas9 edited *miPEP*^*CR*^ lines grown on ½ strength Murashige and Skoog (MS) medium supplemented with water (control) and 0.25 µM miPEP858a. Data are plotted as means ±SD. Error bars represent standard deviation. Asterisks indicate a significant difference between the treatment and the control according to two-tailed Student’s *t*-test using Graphpad prism 5.01 software (n=30 independent seedlings, (* P < 0.1; ** P < 0.01; *** P < 0.001). **(E)** Western blot analysis of CHS protein in 5-day old WT and CRISPR/Cas9 edited *miPEP*^*CR-2*^ line grown on ½ strength Murashige and Skoog (MS) medium (-) and (+) supplemented with 0.25 µM miPEP858a (+). **(F)** Histochemical staining of stem sections of 35-day old WT and *miPEP*^*CR-1*^ with phluoroglucinol was performed after spraying with water (control) and 0.25 µM miPEP858a (scale bars, 200 µm).

**Figure 5.**
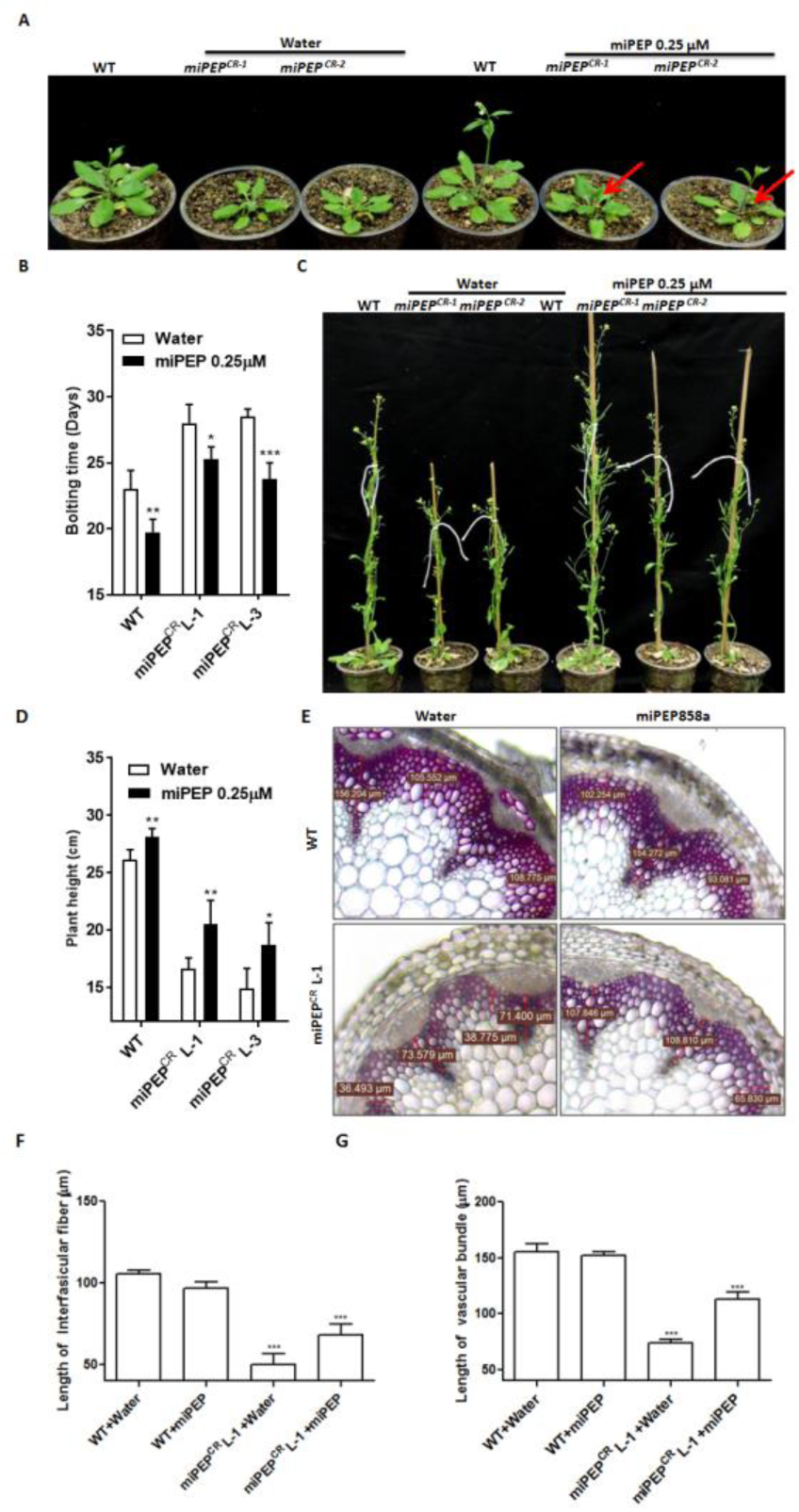
Spraying of miPEP858a leads to complementation of phenotype in CRISPR/Cas9 edited *miPEP*^*CR*^ lines. **(A, B)** Change in bolting time of WT and miPEP CRISPR edited lines after spraying with water and miPEP858a. **(C, D)** Change in plant height (cm) of WT and miPEP CRISPR edited lines after spraying with water and miPEP858a. **(E)** Transverse section of stem of 35-day old WT and miPEP858a CRISPR edited plant sprayed with water and miPEP858a, were stained with phluoroglucinol to show change in lignin content. **(F)** Bar graph showing the length of interfascicular fibers (µm) of WT and miR858CRISPR edited plants sprayed with water and miPEP858a. **(G)** Bar graph showing the length of vascular bundles (µm) of WT and miR858CRISPR edited plants sprayed with water and miPEP858a. Data are plotted as means ±SD.. Error bars represent standard deviation. Asterisks indicate a significant difference between the WT and the edited plants according to two-tailed Student’s t-test using GraphPad Prism 5.01 software (n=5, independent seedlings, * P < 0.1; ** P < 0.01; *** P < 0.001).

**Figure 6.**
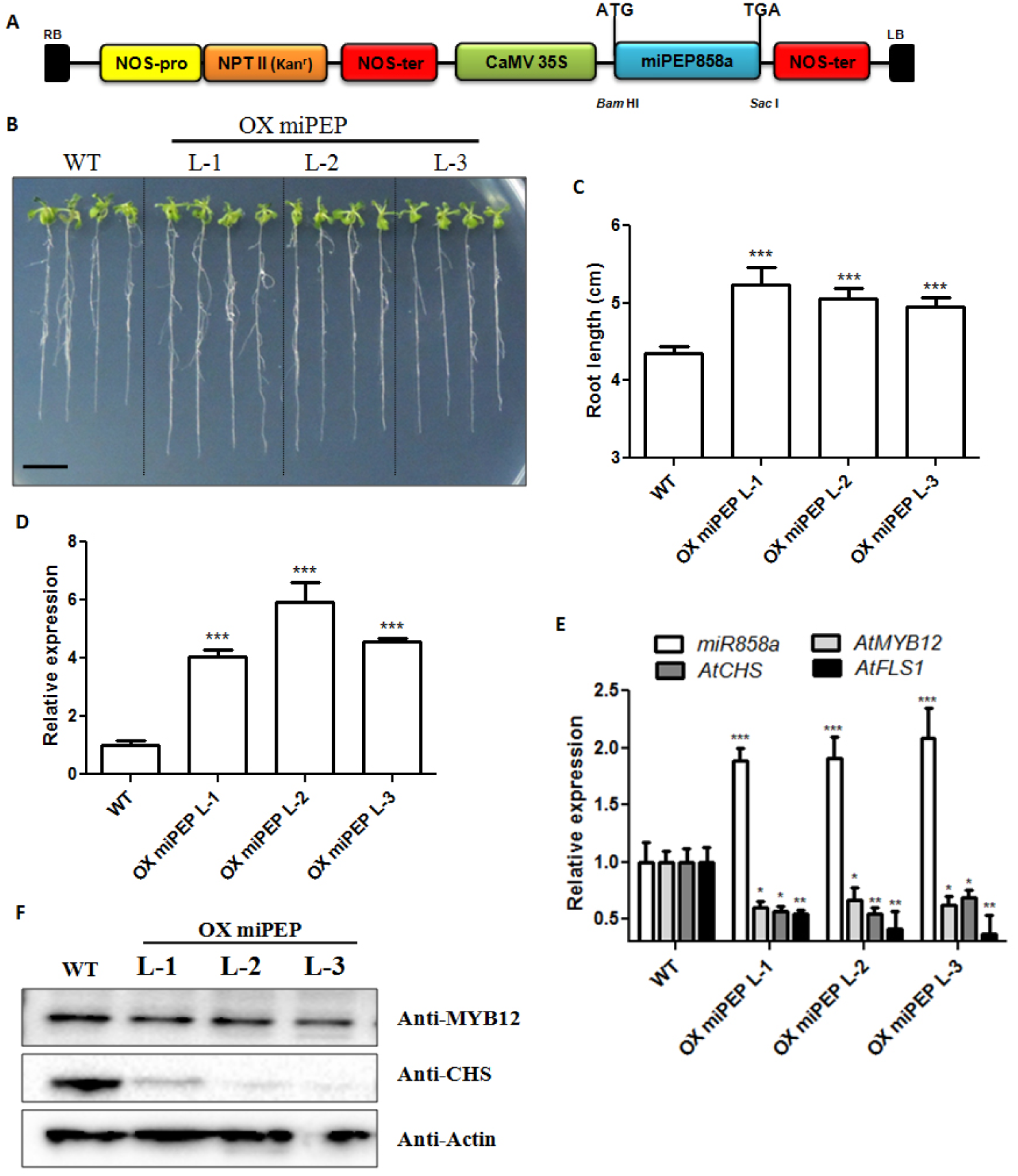
Overexpression of miPEP858a affects plant development and altered expression of flavonoid biosynthesis genes/proteins. **(A)** A schematic representation of OXmiPEP858a construct. **(B)** Representative photographs of the phenotype of 10-day old WT (Col-0) and OXmiPEP seedlings grown on ½ strength Murashige and Skoog. (Scale bars, 1 cm). **(C)** Root length of 10-day old WT (Col-0) and OXmiPEP seedlings grown on ½ strength Murashige and Skoog. Data are plotted as means ±SD. Error bars represent standard deviation. Asterisks indicate a significant difference between the WT and OXmiPEP lines according to two-tailed Student’s *t*-test using GraphPad Prism 5.01 software (n=30 independent seedlings, * P < 0.1; ** P < 0.01; *** P < 0.001). (**D)** Quantification of miPEP858a in WT and OXmiPEP seedlings. (**E)** Quantification of pre-miR858a and target genes in WT and OXmiPEP seedlings. (F) Western blot analysis of MYB12 and CHS protein in 30-day old rosette of WT and OXmiPEP lines. Actin was used as loading control.

**Figure 7.**
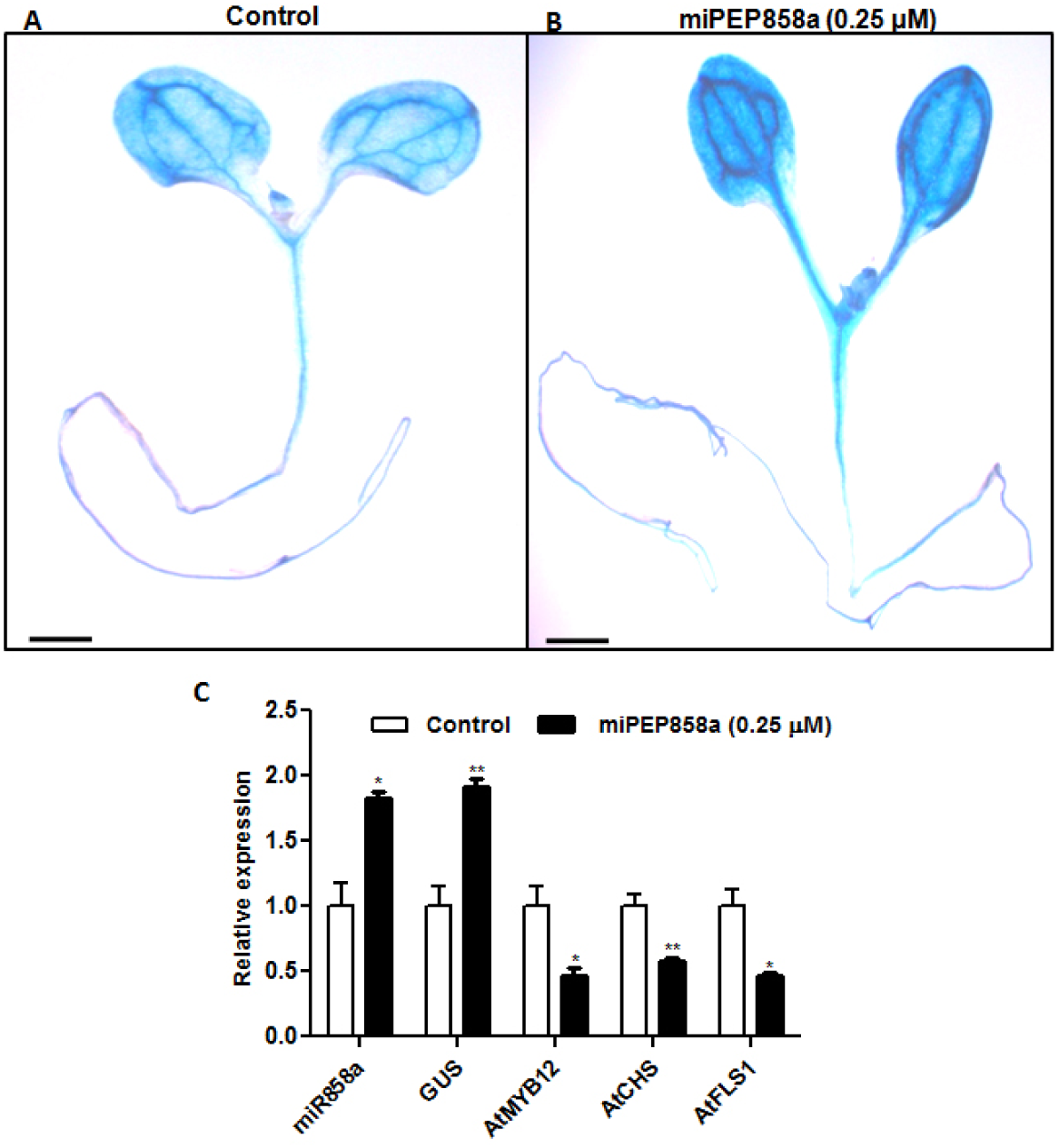
Transcriptional regulation of miR858a is dependent on miPEP858a peptide. **(A, B)** GUS activity in *ProMIR858:GUS* transgenic seedlings grown on ½ strength Murashige and Skoog (MS) medium supplemented with water (control) and 0.25 µM miPEP858a (scale bars, 1 cm). **(C)** Relative expression of pre-miR858a, *GUS, AtMYB12, AtCHS* and *AtFLS1* in *ProMIR858:GUS* expressing seedling grown on ½ strength Murashige and Skoog (MS) medium supplemented with water (control) and 0.25 µM miPEP858a. Data are plotted as means ±SD. Error bars represent standard deviation. Asterisks indicate a significant difference between the treatment and the control according to two-tailed Student’s t-test using Graphpad prism 5.01 software (n=30 independent seedlings, (* P < 0.1; ** P < 0.01; *** P < 0.001).

All the developed mutants displayed a significant decrease in root length as compared to WT on ½ strength MS media (Figure 2A and B). To study whether the mutation in miPEP858a affects the level of miR858a and expression of target genes, qRT-PCR analysis was performed. Expression analysis showed a significant decrease in the expression of miR858a, miR858b and up-regulation of target transcription factor (*AtMYB12*) which regulates expression of structural genes associated with flavonoid biosynthesis (*AtCHS* and *AtFLS1*) (Figure 2C, F and Supplemental Figure 8) (Stracke et al., 2007; Misra et al., 2010; Bhatia et al., 2018). These results are similar to our previous findings, which suggested the importance of miR858a in regulating flavonoid biosynthesis as well as plant growth and development (Sharma et al. 2016).

**Figure 8.**
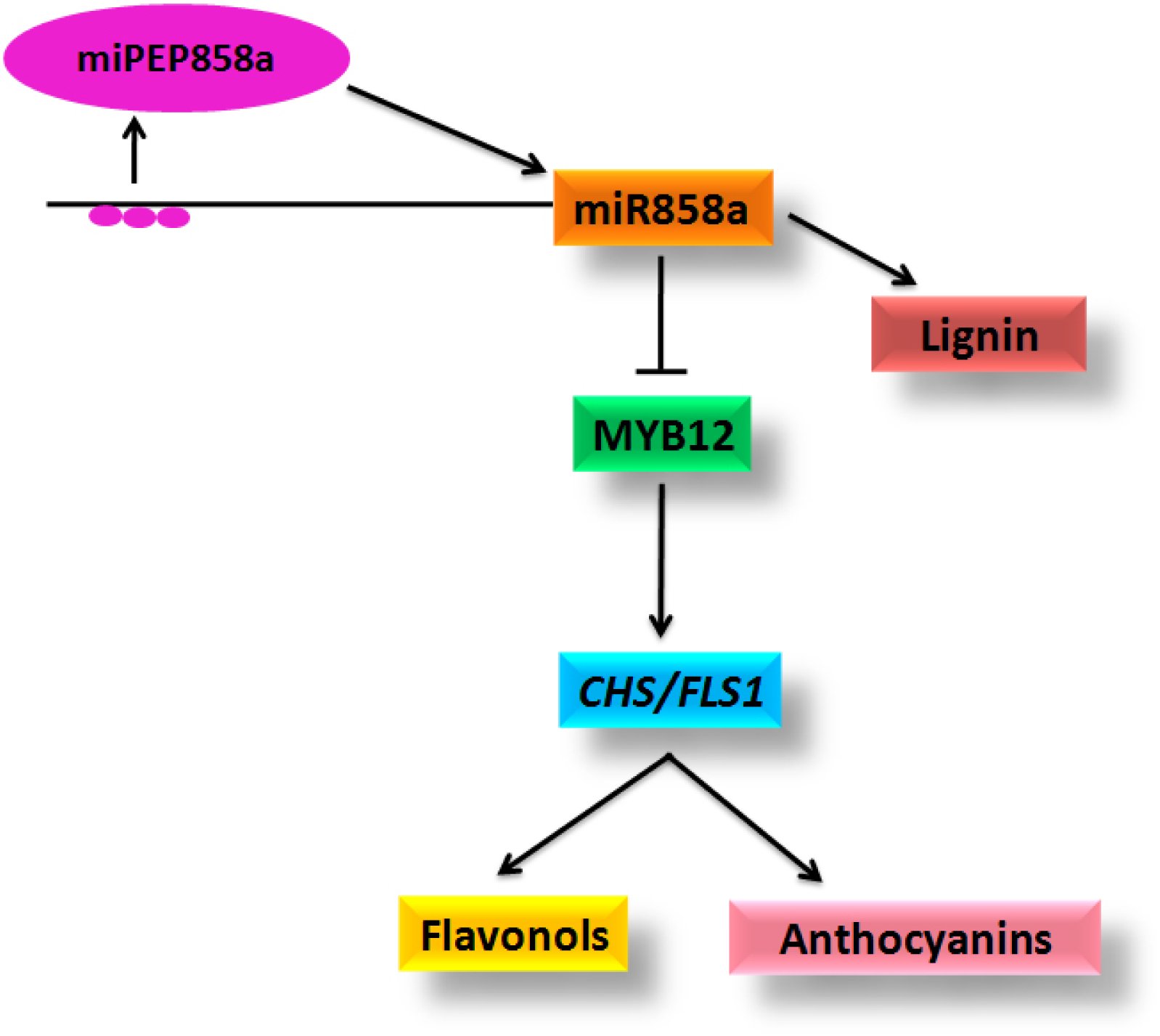
Proposed model for regulation of phenylpropanoid pathway via miR858a and miPEP858a. miPEP858a, short ORF encoded by precursor region of miR858a is involved in transcriptional regulation of miR858a. miR858a negatively controls the levels of MYB12 transcription factor which leads to lesser expression of CHS and FLS1 enzymes thereby causing decreased production of flavonoids. Decrease in flavonoids is accompanied by increase in the lignin content as a result of increase in the flux towards this branch. Arrowheads and tees indicate positive and negative regulation respectively.

Further, to test whether a mutation in miPEP858a has similar effects as mutations in other regions of miR858, plants carrying mutations were grown on soil and were monitored for developmental and phenotypic variations. All the mutants displayed similar phenotypes like delayed bolting, weaker stem, smaller rosette diameter, along with several other developmental variations as compared to WT (Figure 2D and E). These phenotypes were opposite to transgenic plants overexpressing miR858a. To provide further evidence to confirm the role of miPEP858a in regulating miR858 function, AtMYB12 and AtCHS protein levels were analysed using western blot analysis. The analysis suggested a significant increase in the accumulation of MYB12 and its target CHS protein in *miPEP*^*CR*^ plants similar to its accumulation in other *miR858*^*CR*^ plants (Figure 2G). This suggested that *miPEP*^*CR*^ leads to increased abundance of MYB12 protein leading to higher CHS accumulation.

Together, these results suggest that mutations in the miPEP858a coding region, mature miRNA and pre-miRNA confer similar effect on various developmental phenotypes. Results obtained elucidates that miPEP858a controls the phenylpropanoid pathway via regulating the expression of the miR858a which regulates flavonol-specific MYB transcription factors (MYB11, MYB12 and MYB111) in Arabidopsis. Altogether, a significant change in the phenotypes in mutant lines shows that miPEP858a also plays a versatile role in plant growth and development of Arabidopsis.

### Modulated levels of flavonols, anthocyanin and lignin in edited plants

To assess the role of miPEP858a on the synthesis and accumulation of metabolites, we estimated total flavonols, anthocyanin and lignin content in mutant plants. The analysis suggested that *miPEP*^*CR*^,*miR858a*^*CR*^, *miR858b*^*CR*^, *miR858a/b*^*CR*^ and *pre-miR858a*^*CR*^ plants with enhanced expression of target (MYBs) and other related genes leads the redirection of metabolic flux towards the synthesis of flavonols and anthocyanin at the cost of lignin production (Figure 3A and B, Supplemental Figure 9 and 10).

Lignin staining of the cross-section of mutated plants showed decreased lignification in vascular and interfascicular tissues (Figure 3C-F). In order to unravel the molecular mechanism for lesser lignin content in edited plants, the expression analysis of lignin biosynthesis genes including *AtSND1, AtCCR1, AtCAD6* and *AtHCT* was examined by qRT-PCR. Analysis suggested that the expression of these genes is reduced in comparison to WT (Figure 3G). The decrease in expression of these genes is well correlated with the reduction in lignin content of edited plants. Altogether, these results suggest that higher accumulation of flavonoids and anthocyanin in mutated lines is because of increase in expression of miR858a targets genes. Also, the pathway flux is diverted toward flavonoid synthesis at the cost of lignin synthesis.

### Exogenous miPEP858a complements function in *miPEP*^*CR*^ plants

Our analysis suggested that mutation in the miPEP858a coding region causes a decrease in expression of miR858a thereby leading to enhanced expression of the target genes and modulation in the plant growth and development. We, therefore, hypothesized that exogenous application of synthetic miPEP858a to *miPEP*^*CR*^ plants will complement its function. To test this hypothesis, *miPEP858a*^*CR*^ plants were grown on ½ MS with or without supplementation of the synthetic miPEP858a. Supplementation of miPEP858a in media led to the complementation of the root length phenotype of *miPEP*^*CR*^ plants (Figure 4A and B) and increase in the transcription of miR858a with a concomitant decrease in the expression of target and associated genes (*AtMYB12, AtCHS* and *AtFLS1)* (Figure 4C and D). To see the specificity of complementation assay *miPEP858a*^*CR*^ plants were grown on ½ MS with or without supplementation of the NSP (0.25 μM). As expected, supplementation with NSP did not lead to complementation of root length phenotype of *miPEP*^*CR*^ plants (Supplemental Figure 11). In addition, exogenous supplementation of miPEP858a did not complement *miR858*^*CR*^ edited plants (Supplementary Fig. 11). These results suggest that miPEP858a positively regulates expression of miR858a. To strengthen our results, AtCHS protein levels were analysed in WT and *miPEP*^*CR*^ seedlings grown on media supplemented with synthetic miPEP858a. The analysis suggested that supplementation of synthetic miPEP858a to the media leads to the decrease in the CHS protein levels in WT as well as *miPEP*^*CR*^ plants (Figure 4E).

To analyze whether exogenous application of synthetic miPEP858a has a similar effect on plants grown in soil, miPEP858a was sprayed on 14- and 21-day old WT and *miPEP*^*CR*^ plants. After application of exogenous miPEP858a the complete life cycle of these plants was monitored for changes in the phenotype. Application of synthetic miPEP858a on WT and *miPEP*^*CR*^ plants led to early bolting as compared to plants treated with mock (water) (Figure 5A and 5B). On further maturation of these plants, it was found that application of miPEP858a led to significant increase in the plant height in WT as well as *miPEP*^*CR*^ plants as compared to plants treated with mock (Figure 5C and D). Lignin staining and length of interfascicular fibers did not change in WT plants on application of miPEP858a as compared to mock treatment. On the contrary, a significant increase in lignin content and length of interfascicular fibers was observed in *miPEP*^*CR*^ plants on application of miPEP858a. The phenotypes observed on complementation of edited plants with miPEP858a were similar to miR858a overexpression lines (Sharma et al., 2016) (Figure 5E and G). These result suggested that exogenous supply of synthetic miPEP858 to *miPEP*^*CR*^ plants compliments miR858a function not only in seedlings stage but also in mature plants, thereby indicating an important role of miPEP858a in plant growth and development.

### Overexpression of miPEP858a modulates miR858a-associated phenotype

To verify the biological function of miPEP858a, we produced transgenic plants that overexpressed AtmiPEP858a coding region under the control of CaMV35s promoter (Figure 6A). Homozygous transgenic lines which show enhanced levels of miPEP858a were selected for further analysis (Figure 6D). Ten-day old OXmiPEP transgenic lines grown on ½ MS media show enhanced root length as compared to WT (Figure 6B and C).

As expected, the expression of miR858a increased in OXmiPEP lines as compared to WT while decreased expression of target genes (*AtCHS, AtMYB12* and *AtFLS1*) was observed (Figure 6E). Western Blot analysis showed that the levels of AtCHS and AtMYB12 proteins decreased in OXmiPEP lines as compared to WT (Figure 6F). Analysis of metabolites in OXmiPEP lines suggest significantly lower levels of the total anthocyanin and flavonol aglycones (i.e. Kaempferol and Quercetin) levels in OXmiPEP lines as compared to WT. On the contrary, lignin content in OXmiPEP lines increased as compared to WT (Supplemental Figure 13 and 14).The transverse section of stem of OXmiPEP lines showed higher length of vascular bundle and intervascular fiber as compared to WT (Supplemental Figure 13). In addition, expression analysis of genes involved in lignin biosynthesis suggested higher expression of genes in OXmiPEP lines as compared to WT. These results demonstrate the functionality of miPEP858a in controlling miR858a activity.

### Transcriptional regulation of miR858a is dependent on miPEP858a

To explore the possibility that miR858a is regulated in response to miPEP858a peptide, we used promoter lines (*PromiR858a::GUS*) which were developed previously (Sharma et al., 2016). Waterhouse and Hellens (2015) suggested a model and hypothesized that miPEPs might bind to the promoter of respective miRNAs and modulatethe transcription of own genes. To test this hypothesis, we used promoter-reporter lines (*PromiR858a::GUS*) for exogenous peptide assay. Transgenic seedlings were grown on ½ MS with or without supplementation of synthetic miPEP858a (0.25 μM). Interestingly, the expression of *miR858a* and *GUS* genes was significantly higher in plants grown in media supplemented with synthetic miPEP858a as compared to normal control media. Histochemical GUS assay suggested enhanced GUS activity in *PromiR858a::GUS* seedlings supplemented with synthetic miPEP858 in the media as compared to control media (Figure 7A and B). In addition, the expression of target genes (*AtMYB12*) and associated genes (*AtCHS* and *AtFLS1*) decreased significantly in seedlings grown on miPEP858a supplemented media (Figure 7C). These results suggest that miPEP858a regulate its own promoter activity and increases the transcription of the *GUS* gene.

## Discussion

This study provides several lines of evidences to show involvement of miPEP858a in regulation of miR858a expression. Role of miRNAs as agents exercising post-transcriptional control over most eukaryotic genomes is well-known. On the contrary, very limited information is available on how these miRNAs are regulated. In this study, we have investigated transcriptional regulatory mechanism that influences miRNA expression. Earlier, pri-miRNAs were referred to as non-coding RNAs (ncRNAs) as they were thought to be incapable of encoding protein. However, recent studies suggest that a few pri-miRs encode small peptides which participate in regulation of miRNA gene regulation (Lauressegues et al., 2015; Couzigou et al., 2016; Couzigou et al., 2017). In this study, a few peptide coding regions were identified in pri-miRNA sequence of miR858a and functional nature of these peptides was analysed using transient assays in *Nicotiana bethamiana*. Result revealed presence of a sequence encoding small peptide (miPEP858a) of 44 amino acid in pri-miRNA region. A lot of recent studies suggest a role of small peptides as regulators of plant growth and development in plants (Bao et al. 2017; Vanyushin et al. 2017). In corroboration with previous studies, we demonstrate that on the exogenous exposure of miPEP858a to Arabidopsis seedlings increases transcription of pri-miRNA and subsequent down-regulation of target genes of miRNA.

Interestingly, treatment of seedlings with synthetic miPEP858a led to an increase in the root length as a result of enhanced expression of miR858a (Fig. 1C, F and I). In a previous study, authors observed increased nodulation in soybean after treatment with miPEP172c (Couzigou et al., 2017). Collectively, our study suggest that miPEP858a treatment to Arabidopsis plants leads to similar effects as that of miR858a overexpression at both phenotypic and molecular levels (Sharma et al., 2016; Wang et al., 2016). To validate the function of miPEP858a *in planta*, we generated mutant plants defective in members of miR858 family and miPEP858a through CRISPR/Cas9 approach. Recently, single miRNA and miRNA gene families have been targeted via the CRISPR/Cas9 system in Soybean, Arabidopsis and rice (Jacobs et al., 2015, Zhao et al., 2016; Zhou et al., 2017).

Mutation in the coding sequence of miPEP858a reduced the abundance of the pre-miRNA confirming a functional role of this peptide in regulation of miRNA gene expression. Interestingly, blocking miR858a activity by editing miPEP858a resulted in up-regulation of the flavonoid-specific target genes, *AtMYB12*, as well as target genes of *AtMYB12* involved in flavonoid branch of the phenylpropanoid pathway (e.g., *AtCHS1* and A*tFLS1*) (Figure 2E). Estimation of flavonoids (flavonols, anthocyanins) and lignin in Arabidopsis revealed changes in the pattern of flavonoid accumulation between the wild-type and plants with altered expression of members of the miR858 family. In particular, edited plants accumulated higher levels of flavonoids compared with wild-type plants, whereas lignin content reduced in these plants (Figure 3D). Furthermore, complementation of miPEP858a-edited plants with synthetic miPEP858a induced lignin accumulation as compared to mock (water) treated plants (Figure 4F and 5E).

For better understanding of the function of miPEP858a, we produced transgenic lines overexpressing miPEP858a under the control of CaMV35S promoter. The OXmiPEP lines also show gene expression, metabolite levels and phenotype as that of miR858a overexpression lines confirming role of miPEP858a as positive regulator of miR858a activity. Our previous study by Sharma et al., (2016) revealed that *ProMIR858*:GUS activity was constitutive in nature and found in various organs including vascular tissues. We found that lignin content along with length of interfascicular fibers and vascular bundles were higher in OxmiPEP858a plants but lower in miR858a and miPEP858a CRISPR/Cas9 edited plants as compared with the WT. Taken together these results throw light on the impact of miPEP858a on regulating plant growth and development.

Earlier studies suggest specificity of miPEPs towards their associated miRNAs (Lauressergues et al. 2015). Similarly, we observed specificity of the miPEP858a towards regulation of miR858a expression as expression of other miRNAs was not modulated on the application of miPEP858a to Arabidopsis seedlings. Likewise, we also found that use of a non-specific peptide (NSP) does not lead to increase in root length and enhancement in miR858a expression as by miPEP858a. Through promoter-reporter study, our analysis also suggests that miPEP858 regulates own promoter activity as hypothesized by Waterhouse and Hellens (2015). However, uptake of miPEPs by cells, transport to nucleus and interaction with promoter still need detailed investigation (Couzigou et al., 2015).

Collectively, our study proposes a model depicting the importance of miPEP858a in regulating the expression of miR858a thereby exerting its control on the phenylpropanoid pathway leading to flavonoids biosynthesis as well as plant growth and development (Fig. 8). This study added a new layer of regulation of miRNA and confirmed the role of miPEPs in various processes. To provide the further line of evidence, complementation studies performed using synthetic miPEP858a on mutated plants confirmed its vital role in regulating miR858a expression. Our study also suggests that exogenous application of miPEPs can bypass the tedious and complex procedures involved in the production of transgenic plants and can significantly modulate key plant developmental processes with high agronomical importance. It is intriguing to comprehend the molecular mechanism of action of miPEPs in regulation of plant growth and development.

## Acknowledgments

We thank Dr. Qi-Jun Chen, China Agriculture University for pHSE401 vector. The research was supported by Council of Scientific and Industrial Research (CSIR), New Delhi, in the form of NCP project (MLP026). P.K.T. also acknowledges Department of Biotechnology, New Delhi for the financial support in form of projects on pathway engineering, genome editing and TATA Innovation Fellowship. A.S., P.K.B., D.S. and C.B. acknowledge Council of Scientific and

Industrial Research, New Delhi and University Grants Commission(UGC) for Senior Research Fellowship respectively.

## Author Contributions

A.S. and P.K.T. conceived and designed the study. A.S., P.K.B. and C.B. participated in the execution of all the experiments. A.S., P.K.B., C.B., D.S. and P.K.T. interpreted and discussed the data. A.S., P.K.B., C.B., D.S. and P.K.T. wrote the manuscript.

## Material and methods

*Arabidopsis thaliana* (Col-0) was used as wild-type (WT) plant and for editing of miPEP858a and miR858 through CRISPR/Cas9 approach and *Nicotiana benthamiana* was used for transient expression. For germination, seeds were surface sterilized and placed on ½ strength MS medium (Hi-media) containing 1.5% sucrose, pH 5.72-5.8. After stratification for 2 days at 4° C in the dark, plates were transferred to a growth chamber (Conviron) set at 16-h-light/8-h-dark photoperiod cycle, 180 µmol m^-2^ s^-1^ light intensity,22°C temperature and 50-60% relative humidity. Ten-day-old seedlings were used for root length measurement. miR858a overexpressing (miR858OX), MIM858 transgenic lines and *PromiR858a::GUS* transgenic lines were developed earlier (Sharma et al., 2016) and used in this study for analysis.

### Synthetic peptide assay

Synthetic peptides were synthesized through LinkBiotech (http://www.linkbiotech.com).The purity of synthetic peptide is >95%. Peptides were dissolved in water (stock concentration, 5 mM). Seedlings were treated with concentrations from 0.1 to 0.5 μM peptide diluted in the agar medium or in water. For mature plants grown in soilrite, plants were treated with peptide (0.25 μM) diluted in water at 14 and 21 days.

### Peptide sequence

#### miPEP858a

MGGIESLLFTIVRDIGRYGTVCVVYNIKCVYTTRTKASTRTSHP

#### Non-specific peptide

MNIRFSQIAVQDFAKQGSTNISGEFWCSTSQKAYRNTIPKSI

#### miPEP858a Uptake Assays in Arabidopsis Roots

miPEP858a labeled with fluorescent carboxyfluorescein probe (5-FAM) at the N-terminal were purchased from Link Biotech (95–98% purity). 4-day old Arabidopsis WT (Col-0) seedlings were incubated with 50μM 5’fam-miPEP858a-in MG buffer (10mM MgCl_2_ buffer pH 5.8) at 22°C for 12 h. After treatment, seedlings were washed three times by gentle shaking for 5 minutes in MG buffer and roots were imaged using a confocal microscope (Zeiss LSM710 laser-scanning confocal microscope). Fluorescence was visualized as follows: FAM: excitation (ex) 495nm/ emission (em) 545nm.

### Plasmid Construct

For the genome editing using the CRISPR/Cas9 system in *Arabidopsis thaliana*, gRNA for *AtPDS* (AT1G02547) and different regions of miR858a and miR858b were designed using the CRISPR Arizona software (http://www.genome.arizona.edu/crispr). For miR858a, 20 bp gRNA was selected which can target both miR858a and b. For miPEP858a, 20 bp gRNA was selected from the identified coding region. All these gRNA sequences were cloned into binary vector pHSE401 using *Bsa*I restriction site (Xing et al., 2014). This vector contains Cas9 endonuclease-encoding gene under dual CaMV35S promoter as well as genes encoding neomycin phosphotransferase (NPTII) and hygromycin phosphotransferase (HPTII) as selection markers. All the constructs were sequenced from both the orientations using vector forward and vector reverse primer. For OXmiPE858a, ORF was amplified using cDNA of Col-0. Amplicon was cloned into pTZ 57R/T and then transferred into plant expression vector pBI121 containing the cauliflower mosaic virus 35S promoter. For functional ORF analysis, promoter region containing start codon (ATG^1^, ATG^2^ and ATG^3^ and complete ORF^1^) was PCR amplified using high-fidelity enzyme mix (Fermentas) from genomic DNA. Amplicon was cloned into pBI121 plant binary vector using *Hind*III and *BamH*I, restriction site and then transform in Agrobacterium GV3101.

### Transient expression assay

All the constructs were transformed into *Agrobacterium tumefaciens* (strain GV3101) and used to transform in 4 week old *Nicotiana bethamiana* leaves. *A. tumefaciens* strains were resuspended in 5 ml LB (Luria broth) after 24 h incubation at 28°C, 150 rpm secondary inoculation were performed in 50ml (49 ml LB and 1 ml 0.1M MES) culture were incubated 28°C at 150 rpm over night induction medium (10 mM MgCl_2_, 10 mM MES pH5.6, 150 μM acetosyringone) and incubated for 2 h at 28°C. Cultures were then diluted to an A_600_ of 0.5 and injected into *N. benthamiana* leaves using a blunt end 1 ml syringe. Plants were placed under constant light (~70 μmol photons m^2^ sec^1^) for 48–60 h before light measurements, GUS assay were performed in infiltrated leaves.

### Generation of transgenic Arabidopsis plants and analysis of mutations

The pHSE401 vectors having gRNAs and pBI121 vectors were transformed into GV3101, an *Agrobacterium* strain using the freeze-thaw method and *Arabidopsis* (Col-0) plants were used for transformation by using the floral dip method (Clough and Bent, 1998). The collected T_0_ seeds were screened on ½-MS plate containing hygromycin (20 mg/l) for pHSE401 and on kanamycin (50 mg/ml) for pBI121. Positive plants were transferred on soilrite for maturation. Genomic DNA from leaves was isolated using GenElute^TM^Plant genomic DNA miniprep Kit (G2N70-1KT). Target sites were amplified by PCR using primer surrounding target sites. Mutations were detected using Takara Guide-it mutation detection kit (631443) for miR858a, b, pre-miR858a and miPEP858a. For detection of mutation in *AtPDS*, PCR fragments were digested by *Nco*I. PCR products from positive plants were cloned in cloning vector pTZ57R/T and sequenced using M13F and M13R primer. For OXmiPEP858a, cloned sequence was amplified using CaMV35S F and NosT followed by gene-specific primer.

### Histochemical GUS staining

GUS staining was performed using a previously described method (Jefferson et al., 1989). Seedlings of *PromiR858a:GUS* transgenic lines were immersed in a solution containing 100 mM sodium phosphate buffer (pH 7.2), 10 mM EDTA, 0.1% Triton X-100, 2 mM potassium ferricyanide, 2 mM potassium ferrocyanide, and 1 mg ml^-1^ 5-bromo-4-chloro-3-indolyl-b-D-glucuronide at 37°C for 4 h. The chlorophyll was removed by incubation and multiple washes using 70% ethanol. The seedlings were observed under Leica microscope for GUS staining.

### Expression analysis

For qRT-PCR, total RNA was treated with TURBO DNAase (Ambion), and 1 µg was reverse transcribed using the Revert AidH minus First-Strand cDNA Synthesis Kit (Fermentas) as per manufacturer’s instructions. The cDNA was diluted 20 times with nuclease-free water, and 1 µL was used as a template for quantitative PCR performed using Fast SYBR Green mix (Applied Biosystems) in a Fast 7500 Thermal Cycler instrument (Applied Biosystems). For expression analysis, primers from pre-miRNA858a and pre-miR858b regions were used. Expression was normalized using Tubulin and analyzed through the comparative ΔΔ^CT^ method (Livak and Schmitt, 2001). Primer sequences used for expression analysis are listed in Supplemental Table 1.

### Total lignin quantification

To visualize lignified cells in stem, hand-cut sections were stained for 2 min using Toluidine Blue O (Sigma-Aldrich) and 1 min using phloroglucinol blue (Sigma-Aldrich) and visualized on a Leica DM2500 microscope (Kin et al., 2005). Lignin quantification was done by using the method described by Bruce and West (1989). Briefly, samples (approx. 600 mg) were crushed in liquid N_2_ and suspended in ethanol (2 ml). The mixture was centrifugation at 12,000xg (30 min at 4°C). The pellet was dried and resuspended in 5 ml of 2 N HCl and 0.5 ml of thioglycolic acid followed by incubation at 95°C for 8 h and cooling to room temperature. The suspension was centrifuged at 12,000xg for 30 min, and the pellet was washed with double distilled water. After re-centrifuging at 12,000xg and 4°C for 5 min, the resulting pellet was suspended in 5 ml of 1 N NaOH and agitated gently at 25°C for 18 h. After centrifugation at 12,000xg and 4°C for 30 min, 1 ml of HCl was added to the supernatant and the solution was allowed to precipitate at 4°C for overnight. Following centrifugation at 12,000xg and 4°C for 30 min, the pellet was re-dissolved in 3 ml of 1 N NaOH and the lignin was quantified as absorbance at A_280_.

### Total anthocyanin quantification

Total anthocyanin was quantified according to the previously described method (Li et al., 2016). Briefly, seedlings (100 mg) or 30-day-old rosette were crushed in liquid N_2_ and transferred in extraction solution (Propanol:HCl:H_2_O::18:1:81). Samples were heated at 95°C for 3 min followed by incubation at room temperature in the dark for 2 h. After centrifugation, the absorbance of the supernatants was measured at A_535_ and A_650_. Anthocyanin was calculated as (A_535_-2.2A_650_) / g·FW.

### Total flavonols quantification

Ten-day-old seedlings or 30-day-old rosette tissue was extracted in 1 ml of 80% methanol at 4°C for 2 h with shaking. The mixture was centrifuged at 12000xg for 12 min. Supernatants (5 ml) were taken to 2 ml with methanol and subsequently mixed with 0.1ml of aluminium chloride (10% water solution), 0.1 ml potassium acetate (1 M) and 2.8 ml of MQ water. After 30 min incubation, absorbance at 415 nm was recorded. The calibration curve was developed using rutin as the standard. Total flavonol content was calculated as equivalents of rutin used as standard (Loyala et al., 2016).

### Extraction and quantification of flavonols

For extraction of flavonols, seedlings (500 mg) were ground in liquid nitrogen and extracted in 80% methanol overnight at room temperature. The extract was hydrolysed with equal amount of 6 N HCl at 70°C for 40 minutes followed by addition of an equal amount of methanol to prevent the precipitation of the aglycones (Jiang et al. 2015). Extracts were filtered through 0.2 µm filters (Millipore) before analysis by HPLC. All samples were analyzed by HPLC-PDA with a Waters 1525 Binary HPLC Pump system comprising of Water 2998 PDA detector as per method developed by Niranjan et al. (2011).

### Total protein extraction and Western blot analysis

Whole seedlings (100 mg) or 30 day-old rosette tissue were snap frozen in liquid N_2_ and ground in extraction buffer containing [50 mMTris (pH7.5); 150 mM NaCl; 1 mM EDTA; 10% glycerol; 5 mM DTT; 1% protease inhibitor cocktail (Sigma-Aldrich, India); 0.1% Triton-X100]. The homogenate was transferred to a tube and centrifuged at 20,000xg for 12 min at 4°C. The supernatant was transferred to a new tube and aliquot of (5 µl) was used to estimate protein concentration by Bradford assay (Braford, 1976). Laemmli sample buffer (2X) was mixed with protein sample and boiled at 96°C for 5 min. Protein samples (10 µg) were separated on SDS-PAGE (12%) gel at 150 V for 2 h and transferred onto a PVDF membrane at 25V for 2 h in transfer buffer containing Tris-HCl (3.03 g), Glycine (14.4 g), methanol (20%) using Semi-Dry transfer apparatus (Bio-Rad, USA). After transfer, membranes were incubated in the blocking solution (3% non-fat dry milk in Tris-Buffered saline containing 0.05% Tween-20) for 1 h at room temperature. Blots were incubated in primary antibody (diluted in TBS-T) for overnight, followed by washing with wash buffer (Tris Buffered saline containing 0.05% Tween-20) thrice, 5 min each. The secondary antibody, conjugated with horseradish peroxidase diluted (1:10,000 times) in blocking buffer was incubated for 1 h at room temperature followed by washing with wash buffer for 3 times, 5 min each followed by visualisation using ChemiDoc (MP system, Bio-Rad) after incubating with luminol/enhancer and peroxide buffer (1:1 ratio). Commercial antibodies used in the analysis were Anti-Actin Antibody (A0480; Sigma-Aldrich), anti-CHS (aN-20; sc-12620; SantaCruz Biotechnology, www.scbt.com). Polyclonal antibodies were generated in rabbit against a peptide (**CEGSDNNLWHEKENP)** in the MYB12 protein sequence and were affinity purified against the peptide (Eurogenetec).

### Statistical analysis

Data are plotted as means ±SD with error bars as standard deviation. In all the figures, asterisks indicate a significant variation between the WT and the edited plants as per two-tailed Student’s t-test using GraphPad Prism 5.01 software (* P < 0.1; ** P < 0.01; *** P < 0.001). For measurement of root length in Figure 1E, F, G, 2B, 4B (n=30 independent seedlings). For quantification of total anthocyanin, flavonols and lignin content in Figure 3A, B, C (n= 5 independent plants).

## SUPPLEMENTARY INFORMATION

**Supplemental Figure 1.** Upstream sequence (1 kb) from pre-miR858 showing putative ORF. ATG^1^, ATG^2^ and ATG^3^ are start codon of putative ORF. Green highlight sequence showing the functional miPEP858a.Under-lined sequence represents pre-miR858a and mature sequence is shown in blue colour, predicted TSS is gray in colour.

**Supplemental Figure 2.** Specificity of miPEP858a towards miR858a expression. Quantification of various pre-miRNAs in 5-days old WT grown on ½ strength Murashige and Skoog (MS) medium supplemented with water (control) and 0.25 µM miPEP858a. Data are plotted as means ±SD. Error bars represent standard deviation. Asterisks indicate a significant difference between the treatment and the control according to two-tailed Student’s t-test using Graphpad prism 5.01 software (n=30 independent seedlings, (* P < 0.1; ** P < 0.01; *** P < 0.001).

**Supplemental Figure 3.** Analysis of CRISPR/Cas9 edited *AtPDS* mutants. (A) Sequence of gRNA (in red) along with PAM site (in green) showing restriction site of *Nco*I. (B) For mutation analysis, PCR fragments amplified from genomic DNA isolated from Arabidopsis plants transformed with pHSE401+gRNA of *AtPDS.* PCR product was digested with *NcoI* (C) Sequencing analysis of target gene mutations of various lines. Dots indicate deleted bases. Yellow high light denotes the degree of homology between the aligned fragments. Red highlight denotes indel. (D) Seedling and mature plant showing albino phenotype.

**Supplemental Figure 4.** Position of gRNA. pre-miRNA sequence of miR858a and miR858b and ORF of miPEP858a showing location of gRNA in red colour, PAM sequences (NGG) are green in colour.

**Supplemental Figure 5.** Analysis of CRISPR/Cas9 edited miPEP858amutants. (A) For mutation analysis, PCR fragments amplified from genomic DNA isolated from Arabidopsis plants transformed with pHSE401+gRNA of miPEP858a. PCR products were digested with resolvase enzyme (Takara). L-1, L-2 and L-3 are different edited plants. Lanes with or without +R represent PCR product treated or not treated with resolvase enzyme respectively. (B) Sequence analysis of target gene mutations indifferent plants (L-1, L-2 and L-3) as compared to WT. Dots indicate deleted bases. Yellow highlight denotes the degree of homology between the aligned fragments. Red highlight denotes indel. (C) Amino acid sequence of WT and miPEP858a^CR^ edited lines.

**Supplemental Figure 6.** Analysis of CRISPR/Cas9 edited miR858mutants. (A, B, C) represent mutations in miR858a, miR858b and pre-miR858a respectively. Sequence analysis of target gene mutations of various miRNA858 mutant lines is represented in the figure. Mutated lines were selected after digesting with resolvase enzyme (Takara). Dots indicate deleted bases. Yellow highlight denotes the degree of homology between the aligned fragments. Red highlight denotes indel.

**Supplemental Figure 7.** Detection of presence of Cas9 gene in all CRISPR/Cas9 edited lines. PCR detection of Cas9 gene in WT, *miR858a*^*CR*^, *miR858b*^*CR*^, *miR858ab*^*CR*^, *pre-miR858a*^*CR*^ and *miPEP858a*^*CR*^ plants.

**Supplemental Figure 8.** Quantification of pre-miR858b. Quantification of pre-miR858b in 30-day old rosette of WT, *miR858a*^*CR*^, *miR858b*^*CR*^, *miR858ab*^*CR*^, *pre-miR858a*^*CR*^, *miPEP858a*^*CR*^. Data are plotted as means ±SD in the figures. Error bars represent standard deviation. Asterisks indicate a significant difference between the WT and the edited plants according to two-tailed Student’s t-test using Graphpad prism 5.01 software (n=30 independent seedlings, (* P < 0.1; ** P < 0.01; *** P < 0.001).

**Supplemental Figure 9.** Anthocyanin accumulation in CRISPR edited seedlings. Accumulation of anthocyanin in 10-day old WT and CRISPR edited miR858 and miPEP858a seedlings (upper panel). Illustrative picture of the anthocyanin accumulation in WT and CRISPR edited plants (bottom panel).

**Supplemental Figure 10.** Editing of miR858 leads to contrasting accumulation of flavonols (A, Quantification of quercetin and kaempferol content in 5-day old WT, *miR858a*^*CR*^, *miR858b*^*CR*^, *miR858ab*^*CR*^and *pre-miR858a*^*CR*^ plants. Data are plotted as means ±SD. Error bars represent standard deviation. Asterisks indicate a significant difference between the WT and miR858 edited lines according to two-tailed Student’s t-test using GraphpadPrism 5.01 software (n=5, independent seedlings, * P < 0.1; ** P < 0.01; *** P < 0.001).

**Supplemental Figure 11.** Specificity of miPEP858a towards phenotype restoration of CRISPR/Cas9 edited *miPEP*^*CR*^ lines. (A, B) Representative photographs of the phenotype of 10-days old WT and CRISPR/Cas9 edited *miPEP*^*CR*^ lines grown on ½ strength Murashige and Skoog (MS) medium supplemented with water (control), 0.25 µM miPEP858a and 0.25 µM NSP (scale bars, 1 cm). (C) The root length of 10-days old WT and CRISPR/Cas9 edited *miPEP*^*CR*^lines grown on ½ strength Murashige and Skoog (MS) medium supplemented with water (control), 0.25 µM miPEP858a and 0.25 µM NSP. (n=30 seedlings). Data are plotted as means ±SD. Error bars represent standard deviation. Asterisks indicate a significant difference between the treatment and the control according to two-tailed Student’s t-test using Graphpad prism 5.01 software (n=30 independent seedlings, (* P < 0.1; ** P < 0.01; *** P < 0.001).

**Supplemental Figure 12.** Exogenous application of miPEP858a couldn’t leads to restoration of function of CRISPR/Cas9 edited *miR858*^*CR*^ lines. (A, B, C) Representative photographs of the phenotype of 10 days old seedlings of WT and CRISPR/cas9 edited *miR858*^*CR*^ lines grown on ½ strength Murashige and Skoog (MS) medium MS medium supplemented with 0.25 µM miPEP858a and MS medium supplemented with 0.25 µM NSP (scale bars, 1 cm). (D) The root length of 10 days old WT and CRISPR/cas9 edited *miR858*^*CR*^ lines grown on ½ strength Murashige and Skoog (MS) medium supplemented with water (control) and 0.25 µM miPEP858a and 0.25 µM NSP. (n=30 seedlings) Data are plotted as means ±SD in the figures. Error bars represent standard deviation. Asterisks indicate a significant difference between the treatment and the control according to two-tailed Student’s t-test using Graphpad prism 5.01 software (n=30 independent seedlings, (* P < 0.1; ** P < 0.01; *** P < 0.001).

**Supplemental Figure 13.** Overexpression of miPEP858a alters metabolite levels as compared to WT. (A, B, C) Quantification of total anthocyanin flavonols and lignin content in 30-day old rosette and stem of WT and OX miPEP, lines respectively. Data are plotted as means ±SD. Error bars represent standard deviation. (D) Transverse section of stem of 35-day old WT and OXmiPEP lines were stained with phluoroglucinol to show change in lignin content. (E) Bar graph showing the length of interfascicular fibers (µm) of WT and OXmiPEP lines (F) Bar graph showing the length of vascular bundles (µm) of WT and OXmiPEP lines (G) Expression analysis of lignin biosynthesis genes in WT and OXmiPEP lines. Data are plotted as means ±SD. Error bars represent standard deviation. Asterisks indicate a significant difference between the WT and OXmiPEP lines according to two-tailed Student’s t-test using Graphpad Prism 5.01 software (n=5, independent seedlings, * P < 0.1; ** P < 0.01; *** P < 0.001).

**Supplemental Figure 14.** Overexpression of miPEP858a alters flavonoid metabolite levels as compared to WT. (A, B) Quantification of quercetin and kaempferol content in 5-day old WT, OXmiPEP858a plants. Data are plotted as means ±SD. Error bars represent standard deviation. Asterisks indicate a significant difference between the WT and miR858 edited lines according to two-tailed Student’s t-test using GraphpadPrism 5.01 software (n=5, independent seedlings, * P < 0.1; ** P < 0.01; *** P < 0.001).

**Supplementary Table 1.** List of primers used in the study

