## Supplementary information for "miRNA-encoded peptide, miPEP858, regulates plant growth and development in Arabidopsis"

TTCAAATTAGTCAGTGCACAGTTTCATCCCTTTTTTGTATAGGAAATTGTGTTTTTTGTTAACTAAACCTGA
TTTATGTTCGTTATACTAACGCATTCTATCGAATAAATTAATAAAACAAACACACACATACGTAAATAAAA
CAAAAGCACAGACAACATACCGAAGAAAAAGGATTACGGAAGCACATGAATTAACTCATCTCAGAATTA
TTGAAGTTCATCATTGTACTATCAGTAACGGCACAACAGTAACATTCCAAATAAAGGTAAAATATG[3]GTA
CACTGCTTCTCAAAAGGAAATAATAATCGACCACACACACAAAATAAAATAAAACAATATGATTTTTCAA
AACACACACACGAAATGTATTACTTTCACGAAACAATGACAATG[2]TTCATCAGTAGTTTTGTGACATCAGT
AAAAAGCTAGATATCCATATTCTCTGGTGGAACATATTTGGTTATTTCGGAAATTCGTATAGTGGACTTC
GAATTTAAGAGTATAATATGAATTGTCAACATAAGAAATCTAGTTATAACTACGCACATG[1]GGAGGTATC
GAATCTTTACTTTTTACAATAGTGAGAGACATTGGTAGGTACGGTACAGTGTGTGTTGTGTATAATATAA
AGTGTGTATATACCACACGCACGAAGGCAAGCACACGTACGAGTCACCCGTGAGTCAATAAACCAAAGA
GTAATTCGAGTGATTGGGTGACACGTGCAACGTTCTCTACTCCAGATGTTTCATCAAGTTTTGTCGTAAG
CCTTCTATTAACTCATAGAGTTAGTTGTGTGGATTCTCCTCCTCTATTATCTTACATCTTCTTCTATAAGTA
GCTAGGGTTCCTCCTTCACAAAAATATCTTTTTTTCTTTGGTAACCAATAGAATAATAATATTGGGATGGG
TTATATACGGTGGACATCAGAAGATGGAGATGAAGAGTGATTGAATTCGATGCTAGGCTTGACAAAGT
GAGCAATCTTATGTGTTCTCATCTTTGTTTCGTTGTCTGTTCGACCTTGGTCTCGATCTCTCCTCAAAACCCTAATA
CGCGCTTTATCGTTTATTTCATTCATCCATCCTTCTTATGATCATCATAAACTAACGTACATGGTTTATCGGTTTGTTA
TTGATGGGGATTGGGGTAGATAGACGATCGAAACTTTTGATTTCTGAG

**Supplemental Figure 1.** Upstream sequence (1 kb) from pre-miR858 showing putative ORFs. ATG[1], ATG[2] and ATG[3] are start codon of three putative ORF. Green highlight sequence shows the functional miPEP858a.Under-lined sequence represents pre-miR858a and mature sequence is shown in blue colour, predicted TSS is shown in gray colour.

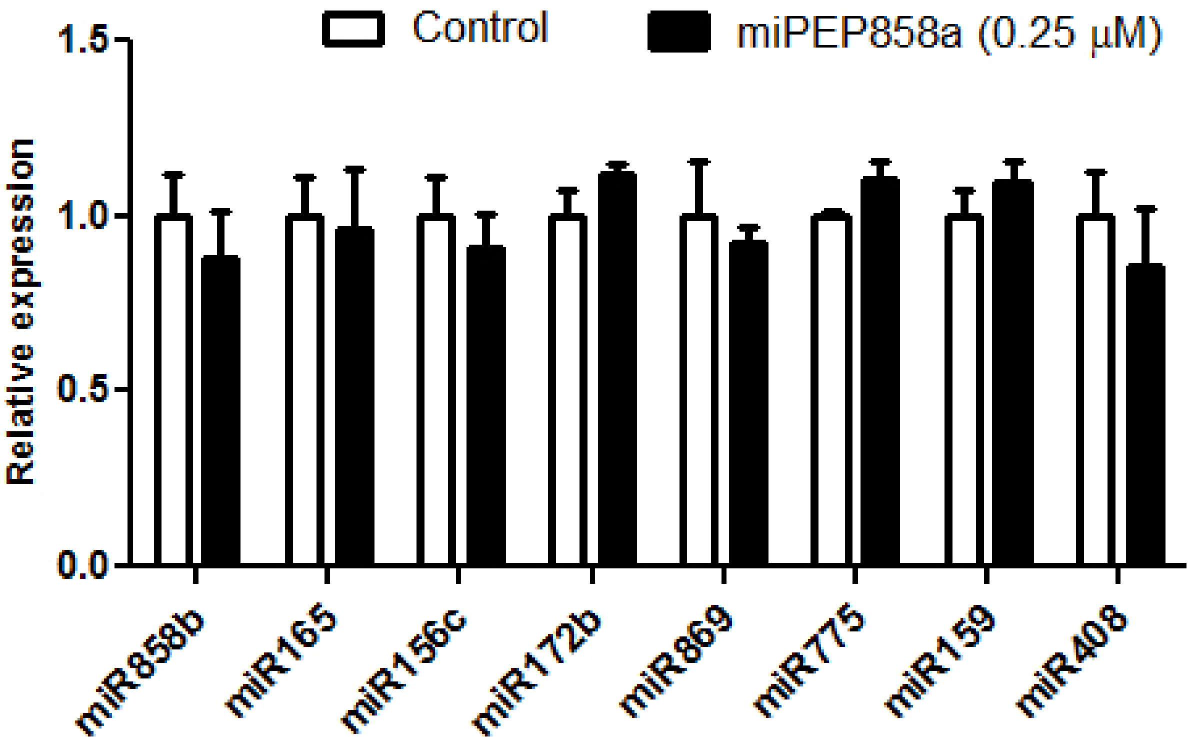


**Supplemental Figure 2.** Specificity of miPEP858a towards miR858a expression**.** Quantification of various pre-miRNAs in 5-days old WT grown on ½ strength Murashige and Skoog (MS) medium supplemented with water (control) and 0.25 µM miPEP858a. Data are plotted as means ±SD. Error bars represent standard deviation. Asterisks indicate a significant difference between the treatment and the control according to two-tailed Student's t-test using Graphpad prism 5.01 software (n=30 independent seedlings, (* P < 0.1; ** P < 0.01; *** P < 0.001).

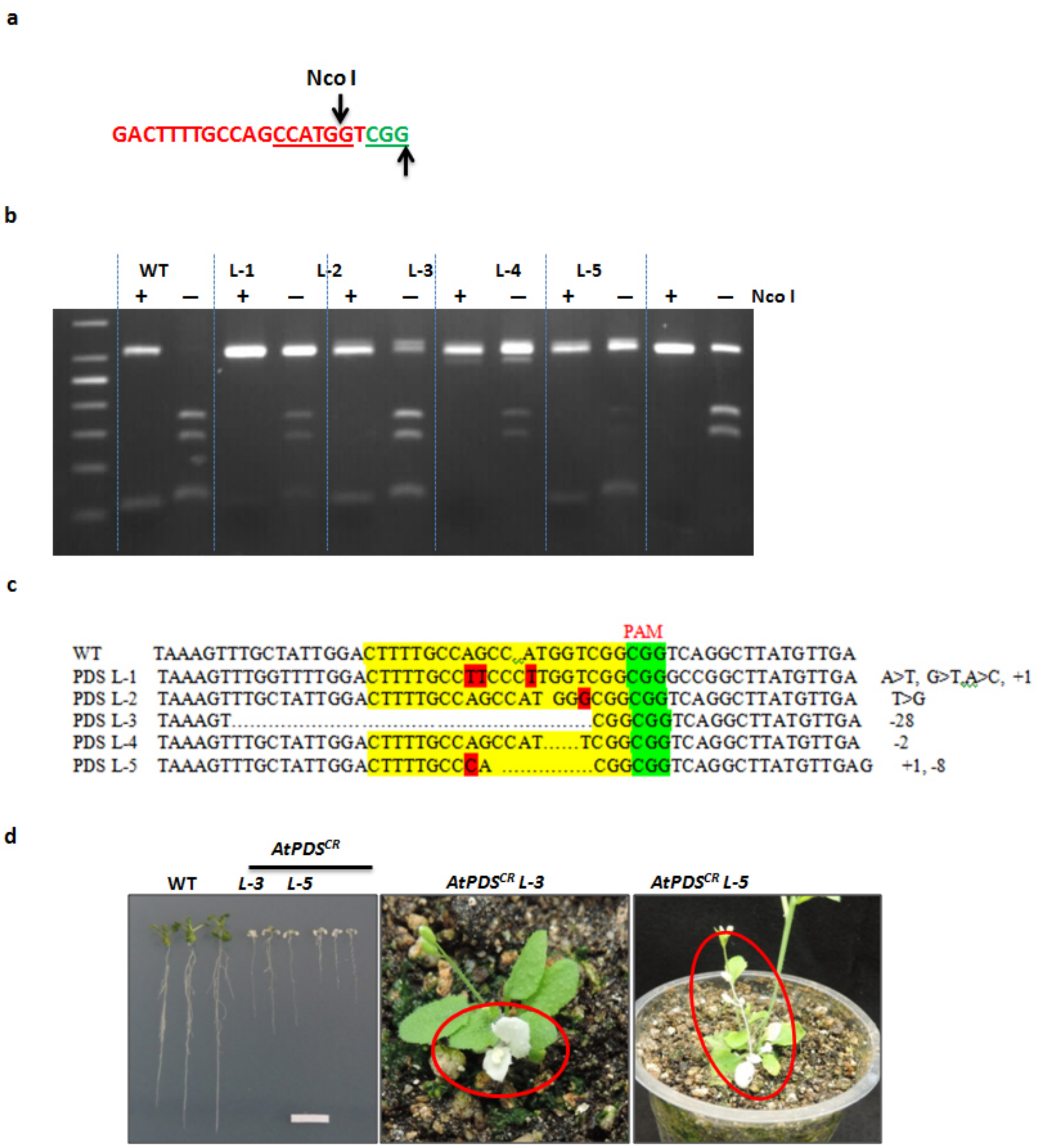


**Supplemental Figure 3.** Analysis of CRISPR/Cas9 edited *AtPDS* mutants. **(a)** Sequence of gRNA (in red) along with PAM site (in green) showing restriction site of *NcoI*. **(b)** For mutation analysis, PCR fragments amplified from genomic DNA isolated from Arabidopsis plants transformed with pHSE401+gRNA of *AtPDS*. PCR product was digested with *NcoI*. **(c)** Sequencing analysis of target gene mutations of various lines. Dots indicate deleted bases. Yellow high light denotes the degree of homology between the aligned fragments. Red highlight denotes indel. **(d)** Seedling and mature plant showing albino phenotype.

**miR858a** **PAM**

CTCATCTTTG**TTTCGTTGTCTGTTCGACCTTGG**TCTCGATCTCTCCTCAAAACCC
TAATACGCGCTTTATCGTTTATTTCATTCATCCATCCTTCTTATGATCATCATAAACTA
ACGTACATGGTTTATCGGTTTGTTATTGATGGGGATTGGGGTAGATAGACGATCGA
AACTTTTGATTTCTGAG

**miR858b**

CCTACCCGAAGGGTTTTGGAGAGTAGACAAAGAGACAAACATAGAGGTGTGAGT
TTGGTTTGGTTTTGGGTTTTGGGGGGTTGGTAGTGTTTGAGGTCCATATGCCTCGA
TCTTCCGCTTCTATTAATTAATTTTCTCTATTAGAAATATTATCTAATTATTCCACTCC
ATCGAACCATGCACATATATAATGATATGTATAAGCCCATCAAAGTTTTTGAATTCA
GTGCAAACACTTGTCAAAGGAGCAGTCCCGATCTGGTTCCATCTTATC**TTCGTTG**
**TCTGTTCGACCTTGG**CTTTGGC
**PAM**

**pre-miR858a**

CTCATCTTTGTTTCGTTGTCTGTTCGACCTTGGTCTCGATCTCTCCTCAAAACCCTA
ATACGCGCTTTATCGTTTATTTCATTCATCCATCCTTCTTATGATCATCATAAACTAAC
GTACAT**GGTTTATCGGTTTGTTATTGTGG**AGGATTGGGGTAGATAGACGATCGA
AACTTTTGATTTCTGA **PAM**

**miPEP858a** **PAM**

**ATG**GGAGGTATCGAATCTTTACTTTTTACAATAGTGAGAGACATTGGTAGGTACG
GTACAGTGTGTGTTGTGTATAATATAAAGTGTGTATATACCACACGCACGAAGGCA
AGCACACGTACGAGTCACCCG**TGA**

**Supplemental Figure 4.** Position of gRNA. pre-miRNA sequence of miR858a and miR858b and ORF of miPEP858a showing location of gRNA in red colour, PAM sequences (NGG) are green in colour.

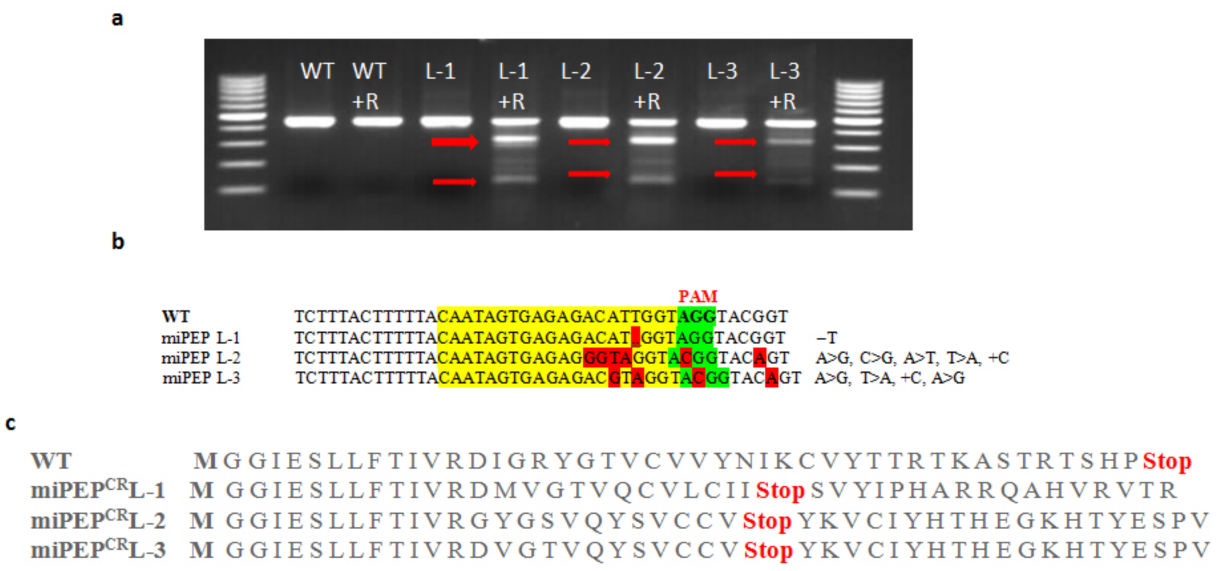


**Supplemental Figure 5.** Analysis of CRISPR/Cas9 edited miPEP858a mutants. **(a)** For mutation analysis, PCR fragments amplified from genomic DNA isolated from Arabidopsis plants transformed with pHSE401+gRNA of miPEP858a. PCR products were digested with resolvase enzyme (Takara). L-1, L-2 and L-3 are different edited plants. Lanes with or without +R represent PCR product treated or not treated with resolvase enzyme respectively. **(b)** Sequence analysis of target gene mutations indifferent plants (L-1, L-2 and L-3) as compared to WT. Dots indicate deleted bases. Yellow highlight denotes the degree of homology between the aligned fragments. Red highlight denotes indel. **(c)** Amino acid sequence of WT and miPEP858a$^{CR}$ edited lines.

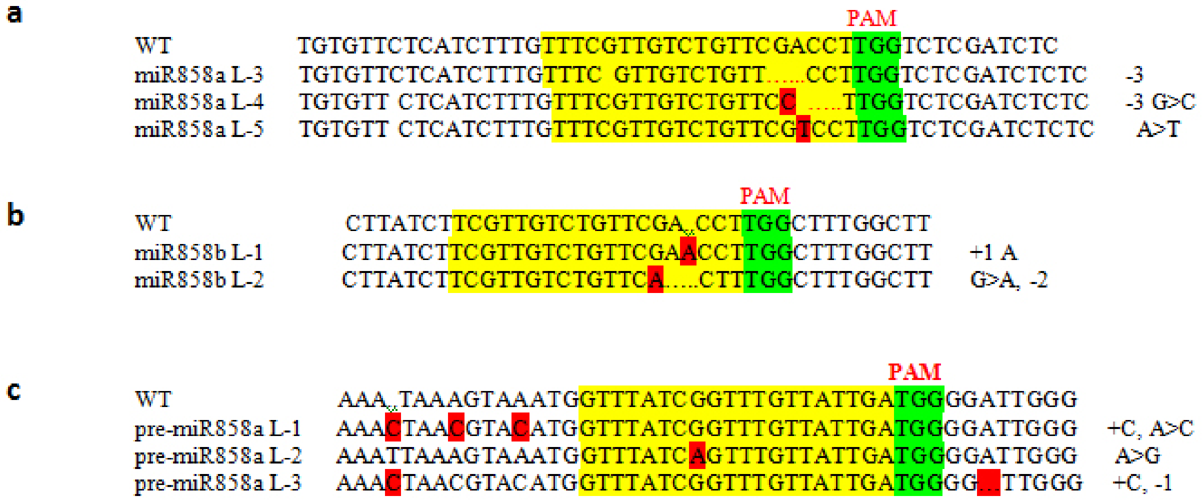


**Supplemental Figure 6.** Analysis of CRISPR/Cas9 edited miR858 mutants. **(a, b, c)** represent mutations in miR858a, miR858b and pre-miR858a respectively. Sequence analysis of target gene mutations of various miRNA858 mutant lines is represented in the figure. Mutated lines were selected after digesting with resolvase enzyme (Takara). Dots indicate deleted bases. Yellow highlight denotes the degree of homology between the aligned fragments. Red highlight denotes indel.

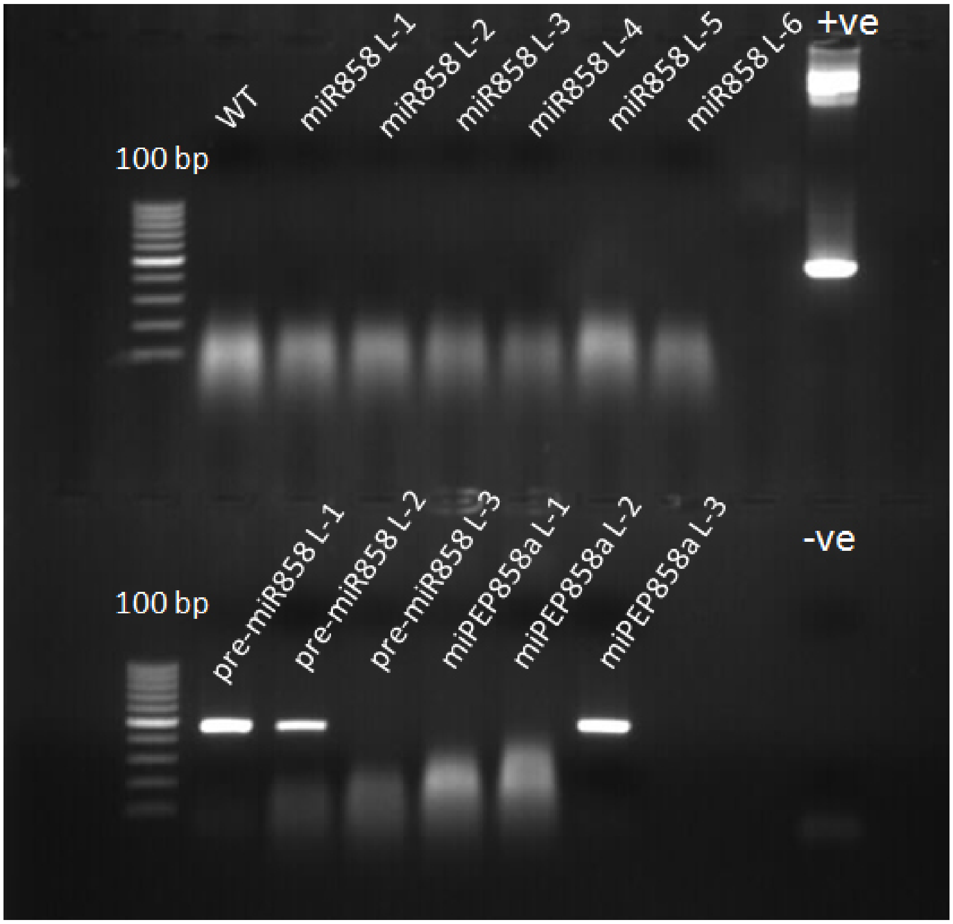


**Supplemental Figure 7.** Detection of presence of Cas9 gene in all CRISPR/Cas9 edited lines. PCR detection of Cas9 gene in WT, *miR858a*$^{CR}$, *miR858b*$^{CR}$, *miR858ab*$^{CR}$, *pre-miR858a*$^{CR}$ and *miPEP858a*$^{CR}$ plants. Analysis suggests that most of lines (except pre-miRNAL-1, pre-miRNAL-2 and miPEP858aL-3) are Cas9-free.

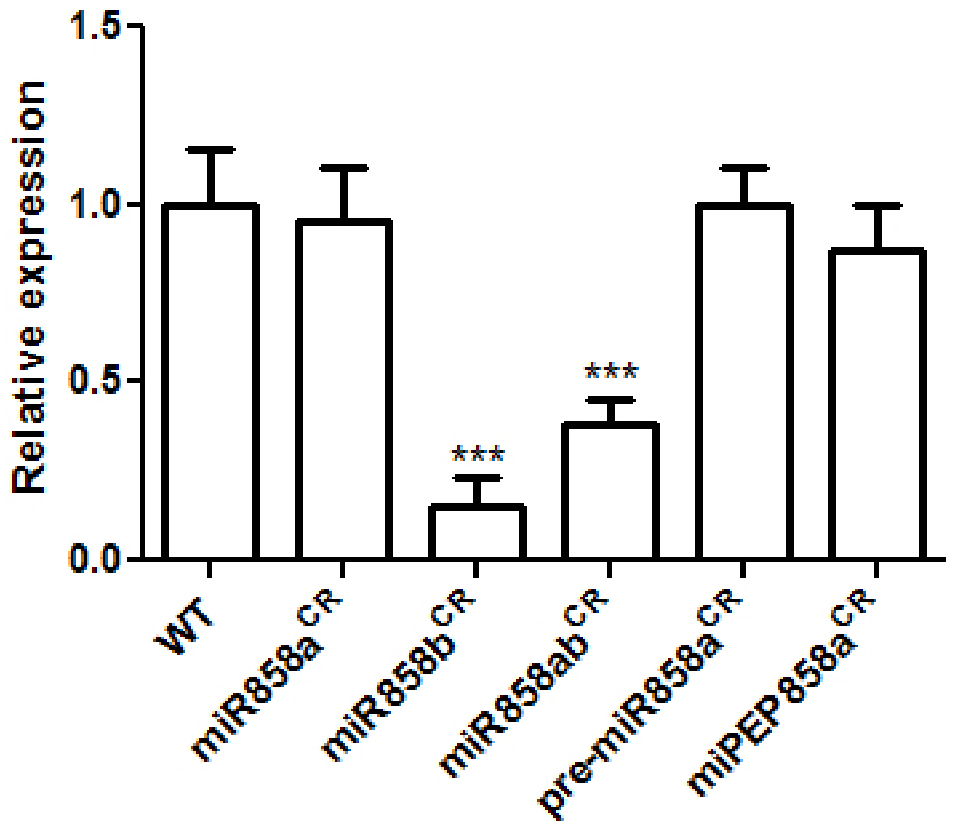


**Supplemental Figure 8.** Expression of pre-miR858b in edited lines**.** Quantification of pre-miR858b in 30-day old rosette of WT, *miR858a*$^{CR}$, *miR858b*$^{CR}$, *miR858ab*$^{CR}$, *pre-miR858a*$^{CR}$, *miPEP858a*$^{CR}$. Data are plotted as means ±SD in the figures. Error bars represent standard deviation. Asterisks indicate a significant difference between the WT and the edited plants according to two-tailed Student's t-test using Graphpad Prism 5.01 software (n=30 independent seedlings, (* P < 0.1; ** P < 0.01; *** P < 0.001).

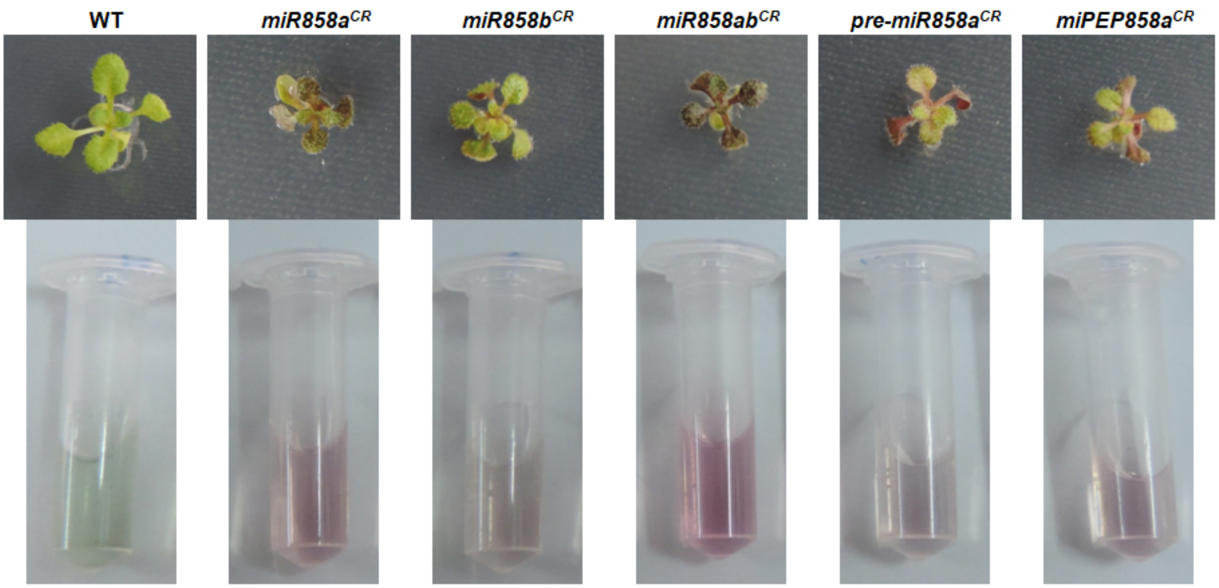


**Supplemental Figure 9.** Anthocyanin accumulation in CRISPR edited seedlings. Accumulation of anthocyanin in 10-day old WT and CRISPR edited miR858 and miPEP858a seedlings (upper panel). Illustrative picture of the anthocyanin accumulationin WT and CRISPR edited plants (bottom panel).

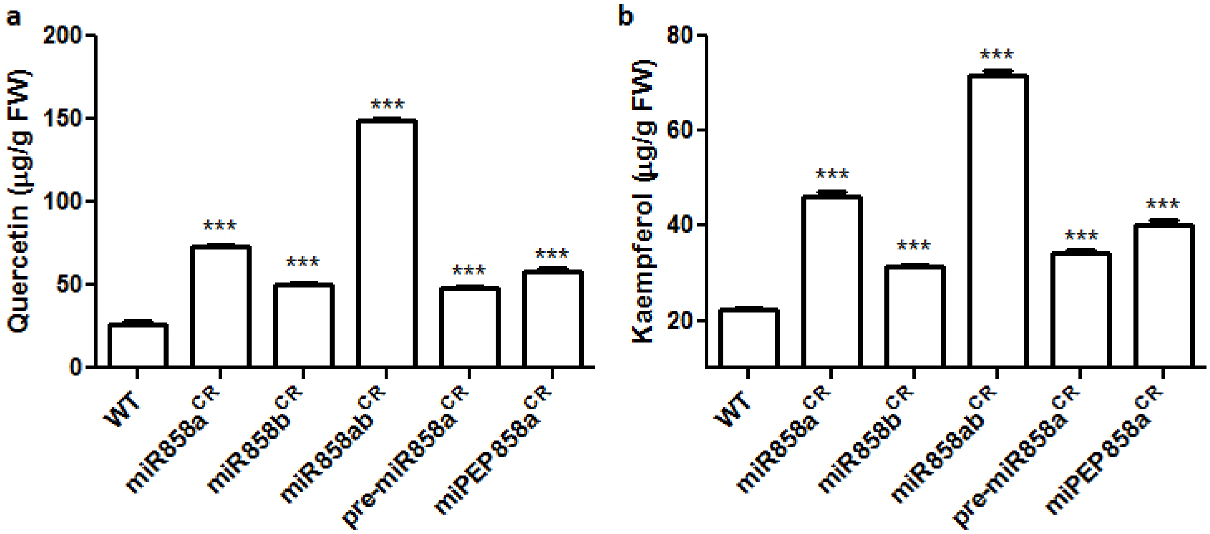


**Supplemental Figure 10.** Editing of miR858 modulates flavonol accumulation. Quantification of quercetin **(a)** and kaempferol **(b)** contents in 5-day old WT, *miR858a*$^{CR}$, *miR858b*$^{CR}$, *miR858ab*$^{CR}$and *pre-miR858a*$^{CR}$ plants. Data are plotted as means ±SD. Error bars represent standard deviation. Asterisks indicate a significant difference between the WT and miR858 edited lines according to two-tailed Student's t-test using Graphpad Prism 5.01 software (n=5, independent seedlings, * P < 0.1; ** P < 0.01; *** P < 0.001).

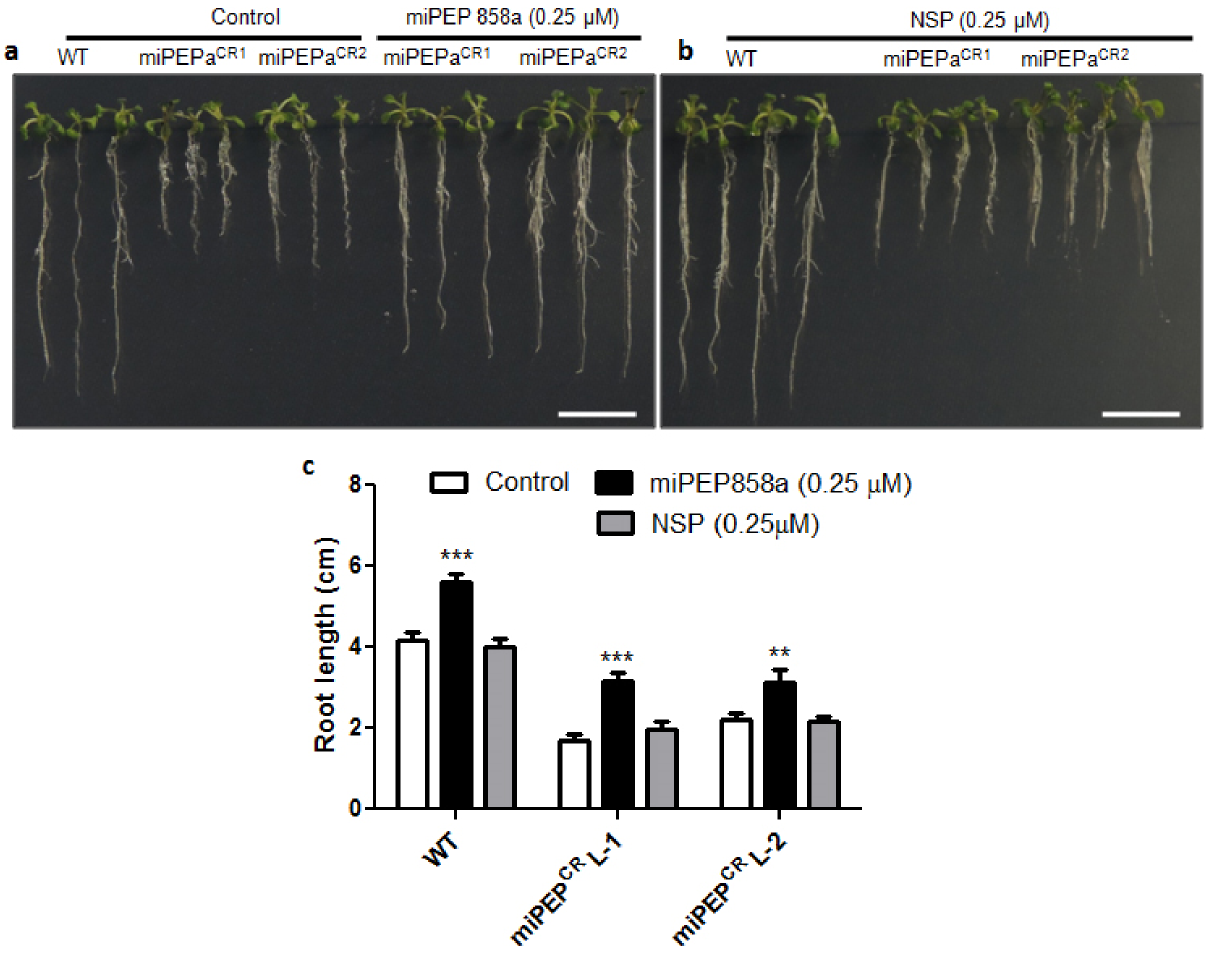


**Supplemental Figure 11.** Specificity of miPEP858a towards phenotype restoration of CRISPR/Cas9 edited *miPEP*$^{CR}$ lines. **(a, b)** Representative photographs of the phenotype of 10-days old WT and CRISPR/Cas9 edited *miPEP*$^{CR}$ lines grown on ½ strength Murashige and Skoog (MS) medium supplemented with water (control), 0.25 µM miPEP858a and 0.25 µM NSP (scale bars, 1 cm). **(c)** The root length of 10-days old WT and CRISPR/Cas9 edited *miPEP*$^{CR}$ lines grown on ½ strength Murashige and Skoog (MS) medium supplemented with water (control), 0.25 µM miPEP858a and 0.25 µM NSP. n=30 seedlings. Data are plotted as means ±SD. Error bars represent standard deviation. Asterisks indicate a significant difference between the treatment and the control according to two-tailed Student's t-test using Graphpad Prism 5.01 software (n=30 independent seedlings, (* $P < 0.1$; ** $P < 0.01$; *** $P < 0.001$).

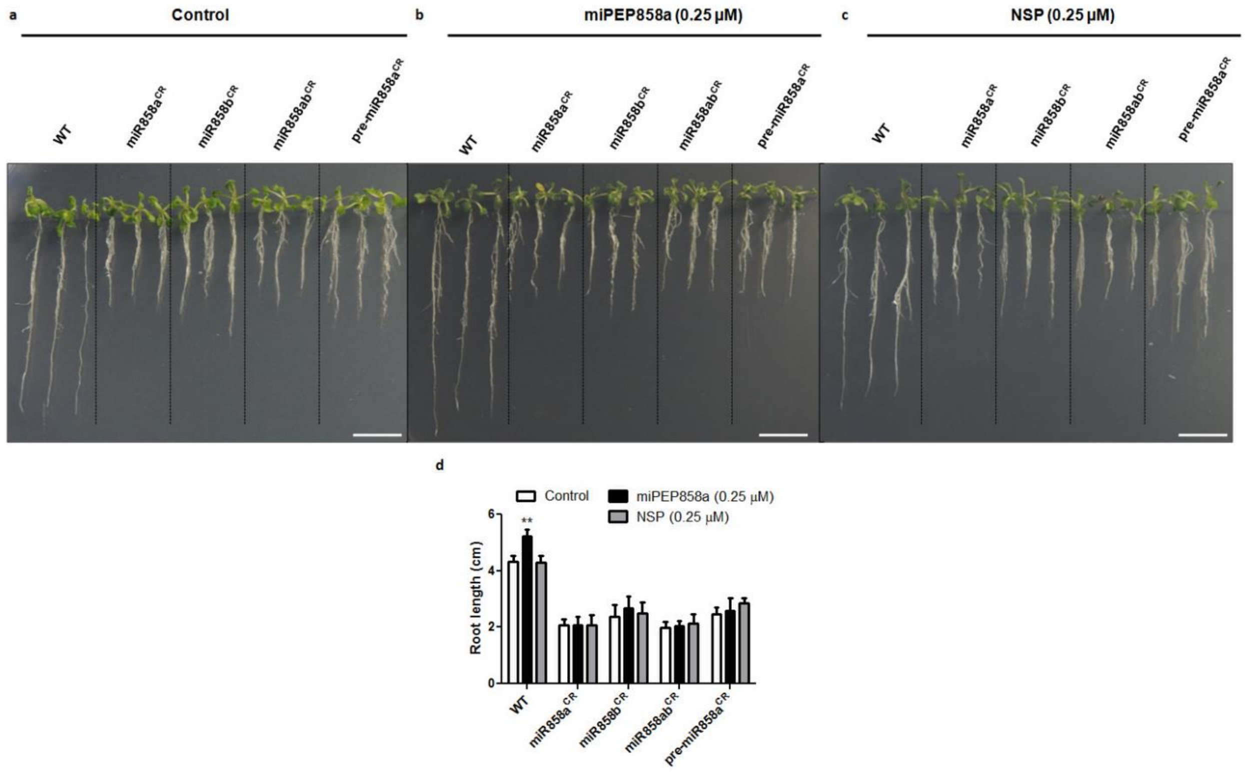


**Supplemental Figure 12.** Exogenous application of miPEP858a and restoration of function of miPEP858a in *miR858*$^{CR}$ lines. **(a-c)** Representative photographs of the phenotype of 10-days old seedlings of WT and CRISPR/cas9 edited *miR858*$^{CR}$ lines grown on ½ strength Murashige and Skoog (MS) medium MS medium supplemented with 0.25 µM miPEP858a and MS medium supplemented with 0.25 µM NSP (scale bars, 1 cm). **(d)** The root length of 10-day old WT and CRISPR/cas9 edited *miR858*$^{CR}$ lines grown on ½ strength Murashige and Skoog (MS) medium supplemented with water (control) and 0.25 µM miPEP858a and 0.25 µM NSP. (n=30 seedlings) Data are plotted as means ±SD in the figures. Error bars represent standard deviation. Asterisks indicate a significant difference between the treatment and the control according to two-tailed Student's t-test using Graphpad Prism 5.01 software (n=30 independent seedlings, (* P < 0.1; ** P < 0.01; *** P < 0.001).

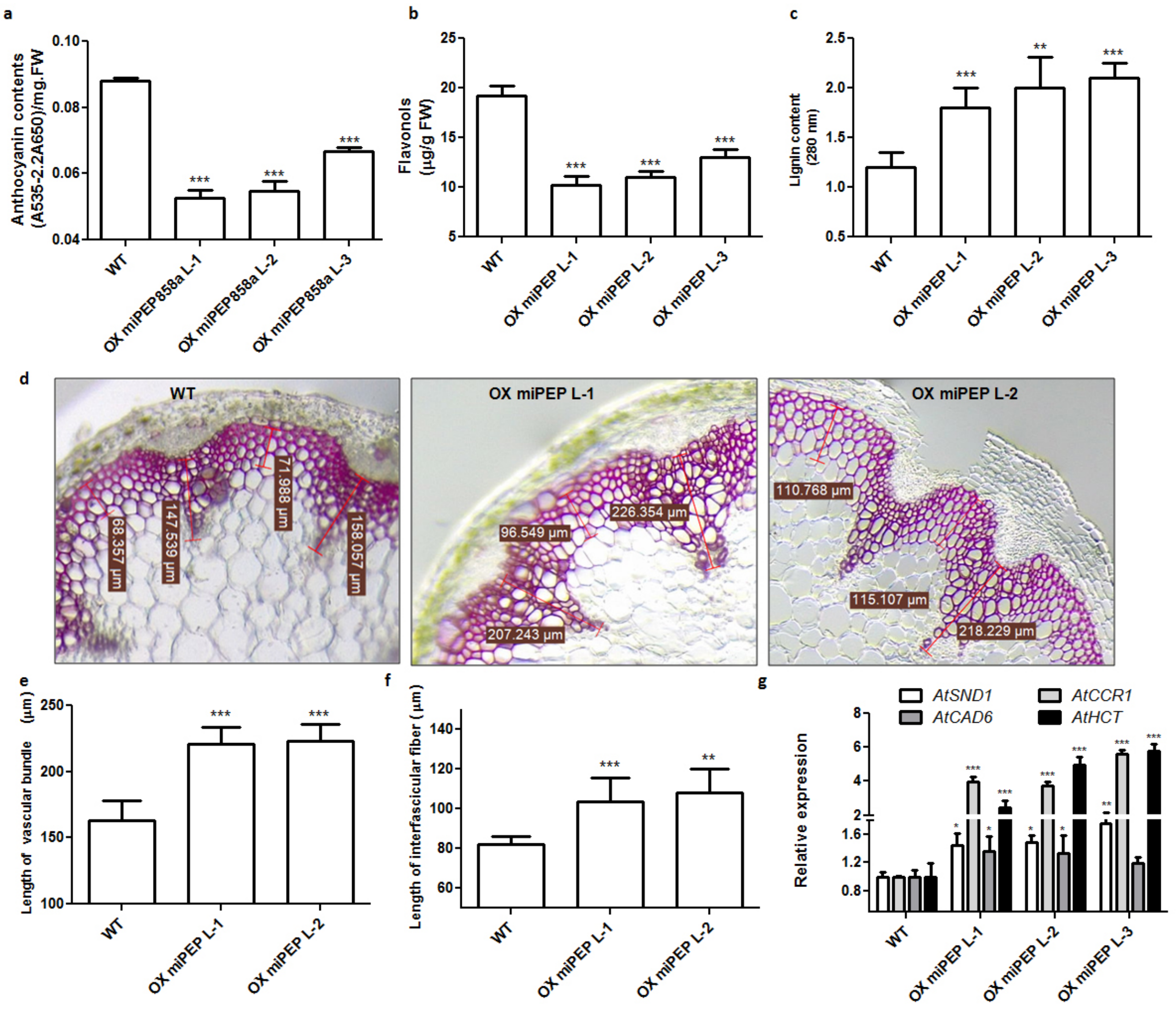


**Supplemental Figure 13.** Overexpression of miPEP858a alters metabolite levels as compared to WT. **(a-c)** Quantification of total anthocyanin flavonols and lignin content in 30-day old rosette and stem of WT and OXmiPEP lines respectively. Data are plotted as means ±SD. Error bars represent standard deviation. **(d)** Transverse section of stem of 35-day old WT and OX miPEP lines were stained with phluoroglucinol to show change in lignin content. **(e)** Bar graph showing the length of interfascicular fibers (µm) of WT and OXmiPEP lines **(f)** Bar graph showing the length of vascular bundles (µm) of WT and OXmiPEP lines **(g)** Expression analysis of lignin biosynthesis genes in WT and OXmiPEP lines. Data are plotted as means ±SD. Error bars represent standard deviation. Asterisks indicate a significant difference between the WT and OXmiPEP lines according to two-tailed Student's t-test using Graphpad Prism 5.01 software (n=5, independent seedlings, * $P < 0.1$; ** $P < 0.01$; *** $P < 0.001$).

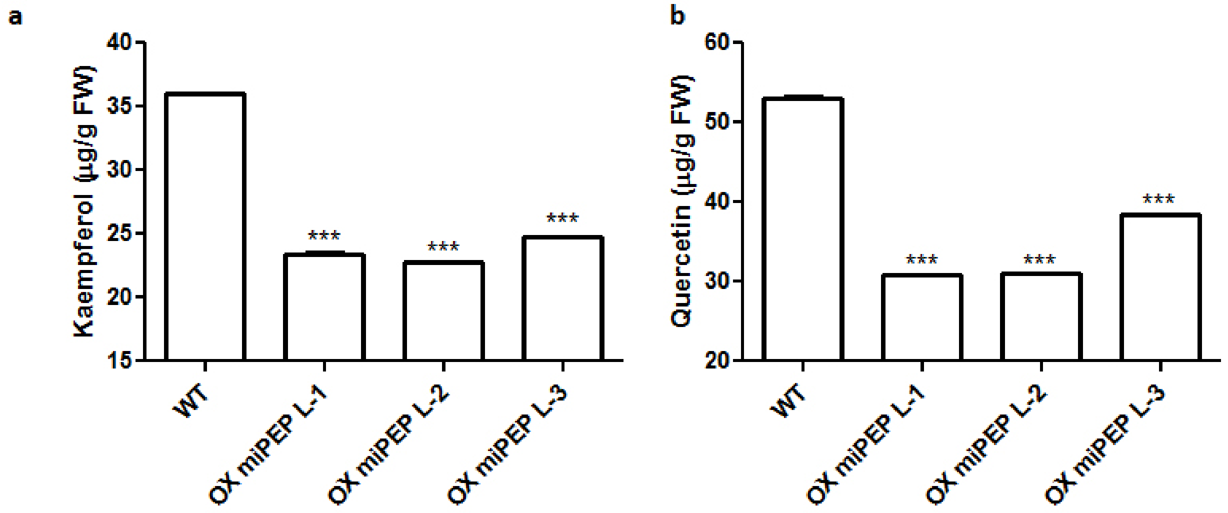


**Supplemental Figure 14.** Overexpression of miPEP858a alters flavonol content as compared to WT. Quantification of quercetin **(a)** and kaempferol **(b)** contents in 5-day old WT, OXmiPEP858a plants. Data are plotted as means ±SD. Error bars represent standard deviation. Asterisks indicate a significant difference between the WT and miR858 edited lines according to two-tailed Student's t-test using Graphpad Prism 5.01 software (n=5, independent seedlings, * P < 0.1; ** P < 0.01; *** P < 0.001).

Supplementary Table 1. List of primers used in the study

| S.No. | Gene | Forward Primer (5′-3′) | Reverse Primer (5′-3′) |
|---|---|---|---|
| 1. | AtPDS gRNA | ATTGGGACTTTTGCCAGCCATGGT | AAACACCATGGCTGGCAAAAGTCC |
| 2. | miR858 gRNA | ATTGTTTCGTTGTCTGTTCGACCT | AAACAGGTCGAACAGACAACGAAA |
| 3. | pre-miR858 gRNA | ATTGGTTTATCGGTTTGTTATTGA | AAACTCAATAACAAACCGATAAAC |
| 4. | miPEP858a gRNA | ATTGCAATAGTGAGAGACATTGGT | AAACACCAATGTCTCTCACTATTG |
| 5. | pHSE401 | TGTCCCAGGATTAGAATGATTAGGC | CCAGAAATTGAACGCCGAAGAAC |
| 6. | At PDS gDNA | AACTGAACTCCGTTGTAGCATTAGCGC | CTACTCTACCAAGTTAAGCTCATCAACATGC |
| 7. | miR858a gDNA | CTTCTTCTATAAGTAGCTAGGGTTCC | CCGTATGTATATACGAACGTAAAGAC |
| 8. | miR858b gDNA | CCCGAAGGGTTTTGGAGAGTAGAC | CGTGAGAGGGTTCCATGTAGCTAAG |
| 9. | pre-miR858 gDNA | CTTCTTCTATAAGTAGCTAGGGTTCC | CCGTATGTATATACGAACGTAAAGAC |
| 10. | miPEPgDNA | AGGTAAAATATGGTACACTGCTTCTC | CCCAATCACTCGAATTACTCTTT |
| 11. | M13 | GTAAAACGACGGCCAGT | CAGGAAACAGCTATGAC |
| 12. | Tubulin | GAGCCTTACAACGCTACTCTGTCTGTC | ACACCAGACATAGTAGCAGAAATCAAG |
| 13. | RT miR858a | CGATCTCTCCTCAAAACCCTA | CGATCGTCTATCTACCCCAATC |
| 14. | RT CHS | GGAGAAGTTCAAGCGCATGTG | ATGTGACGTTTCCGAATTGTCG |
| 15. | RT FLS1 | CCACCGTCATGCGTCAATTACAG | TCTCCGCCGAGACCTTCTTTCAA |
| 16. | RT HCT | GCTCTTAAGGCGAAATCCAAG | CTTTCCCACTGATCTCCACAC |
| 17. | RT miR858b | TGGTTTGGTTTTGGGTTTTG | TGCATGGTTCGATGGAGTGG |
| 18. | RT miPEP858a | GTGAGAGACATTGGTAGGTACGGTAC | GTGATCCGTACGTGTGCTTGC |
| 19. | miPEP EXT | AGGTAAAATATGGTACACTGCTTCTC | CCCAATCACTCGAATTACTCTTT |
| 20. | OX miPEP | ATAAC*GGATCC*AATGGGAGGTATCGAATCT | |
| | TTTGGTTT*GAGCTCT*CACGGGTGACTCGTACGTGCG | | |
| 21. | Ext Pro miPEP858a | TGACCTAAAAGCTTTTCAAATTCAGTG | CCCATCCCAATATTATTATTCTATTGGTTA |
| 22. | Pro miPEP F | CTAA*AAGCTT*TTCAAATTCAGTGGTCTTAC | |
| 23. | ATG[1] | | TAAAGA*GGATCC*ACCTCCCATGTGCGTA |
| 24. | ATG[2] | | GTCACAAA*GGATCC*GATGAACATTGTCATT |
| 25. | ATG[3] | | CCTTTTGA*GGATCC*GTGTACCATATTTTACC |
| 26. | ORF[1] | | ACTCTTTGGTTT*GGATCC*ACACGGGTGACT |
| 27. | RT miR408 | GAGAGAGAGACAGGGAACAAGCA | GAGGCAGTGCATGGGTAGAGAC |
| 28. | RT miR159 | GGCTTTTACTCTTCTTTGGATTGA | CACGCTAAACATTGCTTCGGA |
| 29. | RT miR775 | CGTTGCACTACGTGACATTGAA | CATTGGCACTGCTAGACATCG |
| 30. | RT miR869 | GATGAAGGATTGTGGGATTGTTGC | CCACCACACATGTTTTCATACGG |
| 31. | RT miR165 | CTTGCGCAAAATTACAAAAGC | GCCATGCAAGAAAGATTCAA |
| 32. | pHSE401 Cas9 | CTGACGTAAGGGATGACGCACAA | ATGGAATGCCGATCGGTGTTG |
| 33. | RT miR156c | AGAAGAGAGTGAGCACACAAAGGC | CACTGCTCTATCTGTCAGATTCCG |
| 34. | RT miR172b | AGGCGCAGCACCATTAAGA | GTTGATGCAGCATCATCAAGAT |
| 35. | RT GUS | CCTCGCATTACCCTTACGCTG | CTTGCTGAGTTTCCCCGTTG |
| 36. | RT CCR1 | ACCAAGTGCAAGGACGAGAA | GTCGTAGAGGCTTTGCTTGG |
| 37. | RT SND1 | AAGCTTGAGCCTTGGGATATT | TCCCGGTTGGATACTTCTT |
| 38. | RT CAD6 | TTGGGACGAAAATCGATAGC | TGCTTTTATGCCATGCTCTG |
| 39. | RT MYB12 | TCAGCCGTAAACTCCACAACTT C | CTCAAGACGTCTCCGCCG |

*Italics and under line sequences are restriction sites incorporated in the sequences for cloning.
